# Neuron density fundamentally relates to architecture and connectivity of the primate cerebral cortex

**DOI:** 10.1101/117051

**Authors:** Sarah F. Beul, Claus C. Hilgetag

**Affiliations:** Institute of Computational Neuroscience, University Medical Center Hamburg-Eppendorf, 20246 Hamburg, Germany; Neural Systems Laboratory, Department of Health Sciences, Boston University, 02215 Boston, MA, USA

**Keywords:** anatomical tract tracing, corticocortical connections, connectome, cytoarchitecture, multivariate analyses

## Abstract

Studies of structural brain connectivity have revealed many intriguing features of complex cortical networks. To advance integrative theories of cortical organization, an understanding is required of how connectivity interrelates with other aspects of brain structure. Recent studies have suggested that interareal connectivity may be related to a variety of macroscopic as well as microscopic architectonic features of cortical areas. However, it is unclear how these features are inter-dependent and which of them most strongly and fundamentally relate to structural corticocortical connectivity. Here, we systematically investigated the relation of a range of microscopic and macroscopic architectonic features of cortical organization, namely layer III pyramidal cell soma size, dendritic synapse count, dendritic synapse density and dendritic tree size as well as area neuron density, to multiple properties of cortical connectivity, using a comprehensive, up-to-date structural connectome of the primate brain. Importantly, relationships were investigated by multi-variate analyses to account for the interrelations of features. Of all considered factors, the classical architectonic parameter of neuron density most strongly and consistently related to essential features of cortical connectivity (existence and laminar patterns of projections, area degree), and in conjoint analyses largely abolished effects of cellular morphological features. These results reveal neuron density as a central architectonic indicator of the primate cerebral cortex that is closely related to essential aspects of brain connectivity and is also highly indicative of further features of the architectonic organization of cortical areas such as the considered cellular morphological measures. Our findings integrate several aspects of cortical micro-and macroscopic organization, with implications for cortical development and function.

## Introduction

Corticocortical connections underlie information processing in the cerebral cortex. Recent studies of structural brain connectivity have revealed many intriguing features of complex cortical networks. However, the principles by which connections among cortical areas are organized are still poorly understood. Cortical cytoarchitecture, that is, the cellular composition of the cortical sheet, has been shown to relate to several fundamental aspects of structural connectivity. For example, relationships have been observed between cytoarchitecture and the existence of connections as well as between cytoarchitecture and the laminar patterns of projection origins and terminations in different cortical territories as well as different species (Barbas, 1986; Barbas and Rempel-Clower, 1997; Barbas et al., 2005; Medalla and Barbas, 2006; Hilgetag and Grant, 2010; Barbas, 2015; Beul et al., 2015; Goulas et al., 2016;Hilgetag et al., 2016; Beul et al., 2017). Thus, cortical architecture appears to be a determining factor of the organization of cortical connections. These findings have been integrated into a ‘structural model’ (Barbas, 1986; Barbas, 2015), also called architectonic type principle (ATP), which explains connectivity based on the relative architectonic differentiation of potentially connected areas. Briefly, areas can be categorized along a spectrum of architectonic types, ranging from poorly differentiated types, with low neuron densities and few layers that are hard to demarcate, to highly differentiated types, with numerous, clearly distinguishable layers and high neuron densities. Originally developed qualitatively in the classic studies of Sanides and Pandya (1973), Barbas and coworkers systematically extended the architectonic type principle in quantitative studies across a variety of cortical systems and connection targets in several mammalian species, including prefrontal, parietal, temporal and occipital projection systems, and contralateral as well as subcortical projections (e.g., Barbas, 1986; Barbas and Rempel-Clower, 1997; Rempel-Clower and Barbas, 2000; Dombrowski et al., 2001; Barbas et al., 2005; Medalla and Barbas, 2006; Ghashghaei et al.,2007; Medalla et al., 2007; Hilgetag and Grant, 2010; Goulas et al., 2014; Beul et al., 2015; Goulas et al., 2016;Hilgetag et al., 2016).

However, cortical architecture itself is intricate and varies considerably throughout the cortex, as already detailed in classic studies of brain anatomy (Brodman, 1909;von Economo, 1927; 2009). Moreover diverse aspects of cortical architecture have been measured at different spatial scales. Such measures comprise macroscopic features, such as the laminar appearance of cortical areas, including the thickness of cortical layers and the density and distribution of different types of neurons or glia across layers (Dombrowski et al., 2001; Barbas, 2015). Further macroscopic features are the density of receptors of different neurotransmitter systems (Zilles and Amunts, 2009; Palomero-Gallagher and Zilles, 2017; Zilles and Palomero-Gallagher, 2017) and myeloarchitecture (Nieuwenhuys et al., 2015; Nieuwenhuys and Broere, 2017). In addition, cells within cortical areas have been characterized by a large number of microscopic morphological and physiological measures, such as the density of synaptic spines (Elston et al., 2005; Ballesteros-Yáñez et al., 2006) or firing patterns (Cauli et al., 1997; Dégenètais et al., 2002; Otsuka and Kawaguchi, 2008; Oswald et al., 2013).

Given the abundance of possible features, it remains unclear whether there are aspects of cortical architecture that carry a higher weight in determining corticocortical connectivity than others. Therefore, we analysed the relation of multiple architectonic features to cortical connectivity, to assess the features’ inter-dependence and to identify which of them were most frequently and strongly related to structural cortical connectivity. To be able to evaluate the relative merit of the architectonic features, we devised our analyses such that all features were included conjointly and interrelations between them were taken into consideration, instead of analyzing each measure separately and applying a false discovery rate correction to the significance threshold.

We considered four essential aspects of connectivity, namely the existence and strength of projections, the laminar patterns of projection origins, and the number of connections maintained by an area, the so-called area degree. We probed two groups of measures for their relation to these aspects of structural connectivity. The first group comprised the two macroscopic (area-based) structural features neuron density and spatial proximity. The second group included the four microscopic (cellular) morphological measures soma size, dendritic spine count, dendritic peak spine density and dendritic tree size. Both groups of measures have been individually linked to some features of the macaque connectome in previous reports (e.g., Scholtens et al., 2014; Hilgetag et al., 2016; Beul et al., 2017), but have not yet been combined in a comprehensive analysis that can disclose their comparative relevance.

We considered neuron density because it has been shown to consistently relate to essential aspects of corticocortical connectivity, such as the distribution of projection origins and terminations across cortical layers (i.e., laminar projection patterns), the existence of projections, or topological properties of cortical connectivity (Barbas, 1986; Barbas and Rempel-Clower, 1997; Barbas et al., 2005; Barbas, 2015; Hilgetag et al., 2016; Beul et al., 2017). Neuron density is an objective, quantifiable measure of overall cytoarchitectonic differentiation that is characteristic of different cortices (e.g., Dombrowski et al., 2001).

We also included spatial proximity, not as a measure of cortical architecture, but as an additional macroscopic feature of physically embedded cortical areas that has been shown to be related to connection existence (Young, 1992; Markov et al., 2013a; Beul et al., 2015; 2017) and strength (Douglas and Martin, 2007; Ercsey-Ravasz et al., 2013), but not laminar projection patterns (e.g., Barbas et al., 2005; Hilgetag et al., 2016).

The cellular morphological measures we considered were obtained for pyramidal cells in cortical layer III (L3), based on extensive immunohistochemical analyses (e.g., Elston and Rosa, 1997). The measures comprise the size of the soma, the total spine count of an average pyramidal neuron, the peak dendritic spine density, and the size of the dendritic tree. These measures have been used to quantify ‘pyramidal complexity’ in a previous report that found a relation to topological measures of the macaque connectome (Scholtens et al., 2014). Microscopic, cellular morphological measures appear to be closely correlated with each other, as we also describe below. In primates, neurons show a tendency to become larger, have more complex dendritic arbors and be more spiny towards the frontal cortex (reviewed in Charvet and Finlay, 2014). Characteristics of cellular morphology are crucial cornerstones for how an area can process incoming information. Spine morphology and the spatial arrangement of dendrites directly affect the electrical and biochemical properties of synapses on pyramidal neurons (reviewed in Spruston, 2008; Yuste, 2010), and spine number and density affect the opportunity for neuronal interactions (reviewed in DeFelipe, 2011). These cellular properties, therefore, directly relate to information processing capabilities of cortical populations, especially with regard to the integration of information from numerous sources (Charvet and Finlay, 2014). In line with the areas’ position in the interareal circuitry, morphology in prefrontal association areas allows for a broader integration of inputs (Bianchi et al., 2013; Buckner and Krienen, 2013). Moreover, variation in cellular morphological characteristics across species presumably also reflects differences in the complexity of cortical circuits and specifics of information processing, which plausibly have wide-ranging implications for cognition, memory and learning (DeFelipe, 2011).

Cortical thickness is appealing as a further macroscopic measure because it is relatively easy to measure, also non-invasively. Its relevance to connectivity in healthy and diseased brains has been explored widely (e.g., Lerch et al., 2006; He et al., 2007; Chen et al., 2008; Chen et al., 2011; Tewarie et al., 2014; for a review see Evans, 2013), and it is known to be inversely related to neuron density (von Economo, 1927). However, notwithstanding its appeal as a convenient measure, the usefulness of cortical thickness measures on their own remains open to discussion (Gong et al., 2012) and, rather than individually, thickness measures have recently been used as just one of many measures obtained through MRI to characterize cortical structure (Seidlitz et al., 2018). Moreover, in a direct multivariate comparison with neuron density, we previously reported cortical thickness to be less informative regarding structural connectivity (Beul et al., 2017). Therefore, we did not include cortical thickness in the analyses presented here.

Below, we contrast the relative contribution of each of the six measures (neuron density, spatial proximity, L3 pyramidal cell soma size, dendritic spine count, dendritic peak spine density and dendritic tree size) for characterizing inter-areal connections in the macaque cerebral cortex. Specifically, we considered four essential features of connectivity, projection existence, projection strength, laminar patterns of axon termination, and the number of connections maintained by an area. By employing analyses designed to account for interrelations between the structural measures, we showed that not all measures were equally relevant in predicting connectivity, and that neuron density in particular emerged as the most essential and informative feature for explaining multiple properties of structural cortico-cortical connections. The conclusion is that neuron density constitutes a fundamental architectonic marker of cortical areas that is closely related to diverse macroscopic and microscopic structural cortical features, with principal implications for cortical function and development.

## Materials and Methods

We first introduce the connectome data set of the macaque cortex and then present the investigated structural measures. Subsequently, we describe statistics and procedures used in the analyses.

### Connectivity data

We analysed an extensive, up-to-date set of anatomical tracing data in the macaque cortex (Markov et al., 2014). Based on injections of retrograde tracer in 29 areas, the dataset provided information on 2610 corticocortical connections between 91 areas (represented in the M132 atlas, Markov et al., 2014). That is, projections targeting about a third of the cortex are included in this data set. For projections found to be present, projection strength was given as the fraction of labeled neurons outside of the injected region (FLNe), thus normalizing the number of projection neurons between two areas to the total number of labeled neurons for the respective injection, as done previously (e.g., Barbas and Rempel-Clower, 1997; Medalla and Barbas, 2006). In the data set, projections were included as present without a threshold on projection strength, that is, even a single labelled axon was considered to constitute a present projection. Most cortical areas were injected only once, but controls for consistency between repeated injections were performed in a few areas.

To assess the relation of the structural measures with projection existence, we transformed projection strength to a binary measure (absent vs. present).

To assess the relation of the structural measures with projection strength, we considered the natural logarithm, ln(FLNe). The use of a logarithmic scale was indicated, since the most extreme FLNe value was more than three standard deviations above the mean FLNe value (Buzsáki and Mizuseki, 2014). Moreover, we considered ranked projection strength, to account for the fact that areas that receive many afferent projections might have smaller FLNe values per projection than areas that receive only few afferents. To circumvent this possible bias whereby projections from areas with many afferents could artificially be considered too weak through the normalization by the number of neurons labelled per injection, we ranked afferent projections per target area. Specifically, the strongest projection that targeted each of the injected areas (since retrograde tracers were used, only injected areas receive projections in the used data set) was ranked highest (as rank 1), and successively weaker projections were ranked accordingly (by increasing rank number), up to the number of afferents, which differed between areas. This ranking made projection strength dependent only on the relative strength of a given projection to projections from the same injection. Thus, ranking projections by strength allowed us to assess how projection strength related to the structural measures independent of the precise strength of a projection in terms of the number of neurons and the number of afferents an injected area received, alleviating any distortions that may be caused by these factors.

In addition, we analyzed the laminar patterns of projection origins for this data set (which were published separately from the main part of the data set, in Chaudhuri et al., 2015). The fraction of labeled neurons originating in supragranular layers (*N*_SG_%) was provided for 1602 present projections from all injected areas. Specifically, for each projection, *N*_SG_% was computed as the number of neurons labeled in supragranular layers divided by the sum of neurons labeled in supragranular and infragranular layers. To relate *N*_SG_% to the undirected measure of geodesic distance, we also transformed it to an undirected measure of inequality in laminar patterns, |*N*_SG_%|, where |*N*_SG_%| = |*N*_SG_% - 50| * 2 (cf. Hilgetag and Grant, 2010; Beul et al., 2015). Values of *N*_SG_% around 0% and 100%, thus, translated to larger values of |*N*_SG_%|, indicating a more pronounced inequality in the distribution of origins of projection neurons between infra-and supragranular layers and hence deviation from a columnar (bilaminar) pattern of projection origins. We based our analyses regarding *N*_SG_% on the subset of 1132 projections comprising more than 20 neurons (neuron numbers for each projection are provided in Markov et al., 2014). Thus, we excluded very sparse projections for which assessment of the distribution of projection neurons in cortical layers was not considered reliable, as was done previously (cf. Barbas et al., 2005; Beul et al., 2017). Note that sparse projections were only excluded from analyses involving *N*_SG_%, but not from analyses considering binary projection existence.

The data set contained the 29×29 subgraph of injected areas for which all possible connections have been examined. To assess overall area degree, we considered only areas within this edge-complete subgraph, computing the overall degree of each area as the sum of the number of its efferent and afferent projections, as reported previously (Beul et al., 2017). Separately, we also considered the number of efferent projections, out-degree, and the number of afferent projections, in-degree. In-degree was computed both within the 29×29 edge-complete subgraph, and cortex-wide, using all reported projections between the 91 cortical areas (i.e., on the 91×29 graph).. In two previous publications (Beul et al., 2015; Beul et al., 2017), we investigated the architectonic difference between rich-club and periphery nodes. We found a difference in architectonic differentiation between the two sets of nodes, which however generalized to the relation between architectonic differentiation and area degree that we also analyze here. Therefore, we decided to perform no further analyses of rich-club versus periphery nodes. Note that degree was the only connectivity measure that is a property of cortical areas, rather than corticocortical projections. Therefore, a smaller number of data points was considered here than in other analyses that assessed properties of projections.

### Structural measures

Next, we introduce the investigated six area-based and cellular morphological measures, grouped by their spatial (macroscopic or microscopic) scale (Figure 1).

**Figure 1.**
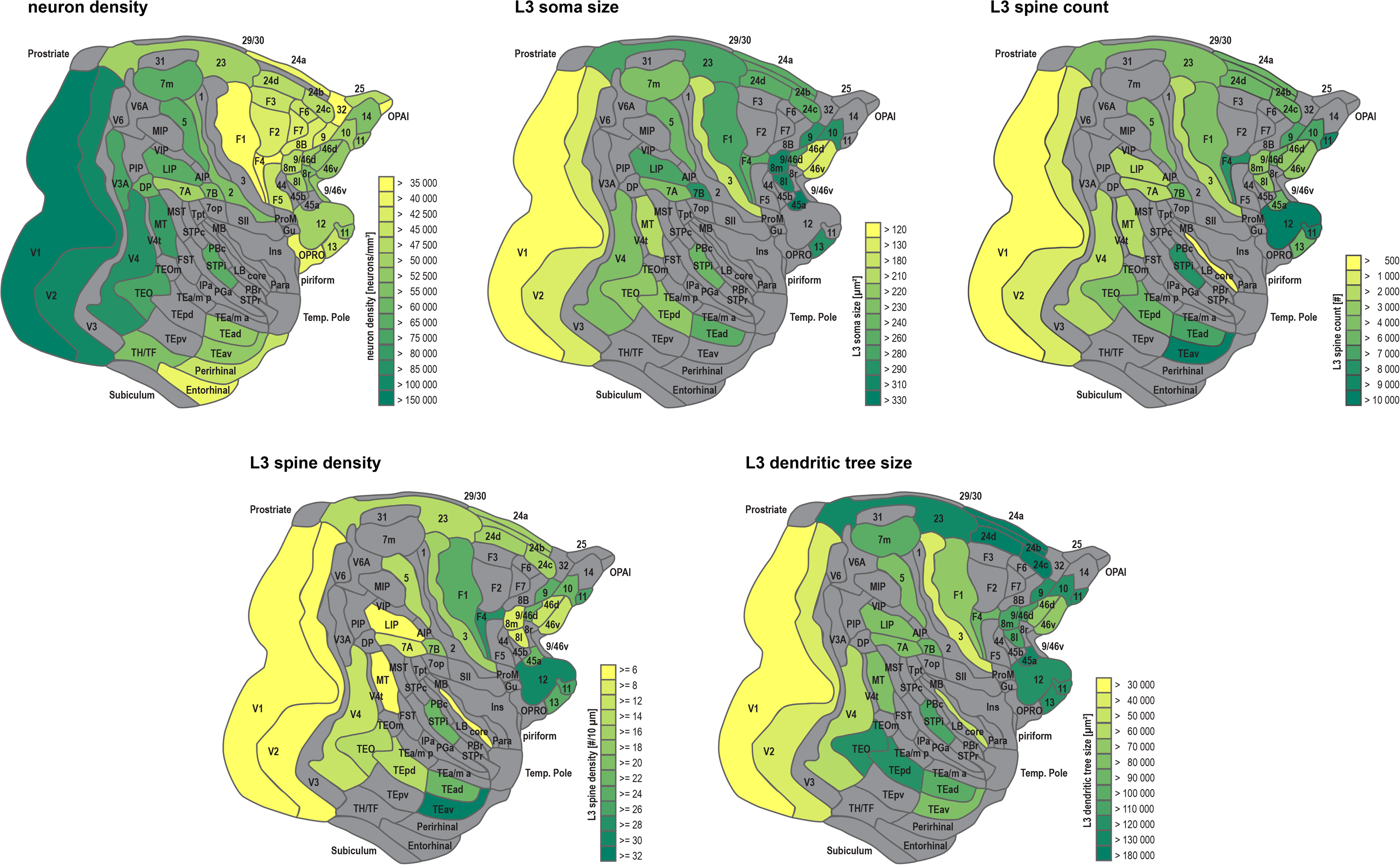
Variation of cytoarchitectonic features across the primate cortex. Variation of neuron density, L3 neuron soma size, L3 dendritic spine count, L3 dendritic spine density and L3 dendritic tree size depicted on the M132 parcellation (Markov et al., 2014). For grey areas, no values were available. See Supplementary Table 1 for correspondences between areas in the M132 parcellation and alternative parcellations. Abbreviations as in Markov et al. (2014).

### Macroscopic measures

#### Neuron density

We used an unbiased quantitative stereologic approach to estimate neuron density from immunohistochemically stained sections of macaque cortex using a microscope-computer interface (StereoInvestigator, MicroBrightField Inc., Williston, VT). The neuron density measurements used here have been partly published before (Dombrowski et al., 2001; Hilgetag et al., 2016; Beul et al., 2017). More detailed descriptions of the acquisition of the neuron density measurements have been published elsewhere (Hilgetag et al., 2016; Beul et al., 2017).In total, neuron density measures were available for 48 of the 91 areas in the M132 parcellation. We quantified how similar areas were in their neuron density by computing the log-ratio of neuron density values for each pair of areas (which is equivalent to the difference of the logarithms of the area densities). Specifically, log-ratio _density_ = ln(density _source area_ / density _target area_). This procedure enabled us to directly relate each sampled projection to the density ratio of its source-and target-area. The use of a logarithmic scale was indicated, since the most extreme value of the neuron density measures was more than three standard deviations above the mean of the considered neuron densities (Buzsáki and Mizuseki, 2014). For analyses which required considering an undirected equivalent of the actual neuron density ratio, we used the absolute value of the log-ratio, |log-ratio _density_|. To relate the neuron density ratio to ranked projections strength, we also ranked the absolute value of the log-ratio of neuron density, |log-ratio _density_|, separately per target area. That is, the smallest absolute neuron density ratio was ranked highest (as rank 1) for each injected area, and successively larger absolute neuron density ratios were ranked accordingly (increasing rank number). Hence, areas of similar neuron density (small absolute ratio) were ranked higher than areas of strongly diverging neuron density (large absolute ratio) relative to each of the injected areas.

#### Spatial proximity

To quantify the spatial proximity of cortical areas, we used the geodesic distance between the mass centers of all 91 areas, which are provided as supplementary material with the study of Markov and colleagues (Markov et al., 2013a). To relate the spatial proximity to ranked projections strength, we ranked geodesic distance separately per target area, analogous to the ranking described for the absolute neuron density ratio above. That is, the smallest distance was ranked highest (rank 1), and successively larger distances were ranked accordingly (increasing rank number).

### Cellular morphological measures

Measures of cellular morphology were mostly reported by Elston and colleagues (Elston and Rosa, 1997; Elston and Rosa, 1998a; Elston and Rosa, 1998b; Elston et al., 1999a; Elston et al., 1999b; Elston, 2000; Elston et al., 2001; Elston and Rockland, 2002; Elston et al., 2005; Elston et al., 2009; Elston et al., 2010a; Elston et al., 2010b; Elston et al., 2011a; Elston et al., 2011b; Coskren et al., 2015; Gilman et al., 2017). Specifically, four aspects of L3 pyramidal neuron morphology were measured across the macaque cortex: the size of the cell soma (soma size), the average total spine count on the basal dendritic tree (spine count), the peak density of dendritic spines (spine density), and the size of the basal dendritic tree (dendritic tree size). Spine density was measured as the number of spines per 10 μm dendrite segment, and peak spine density was then calculated as the average density along the five consecutive 10 μm segments that yielded the highest spine density (see e.g. Elston and Rosa, 1998a). Supplementary Table 1 gives an overview of the correspondence between the parcellations used in the morphological data references and the M132 parcellation, as well as the relevant reports. Specifically, in the M132 parcellation, soma size was available for 30 areas, spine count and spine density for 33 areas, and dendritic tree size for 34 areas.

To quantify how similar areas were in these four morphological measures across the cortex, we computed the difference of their values for each pair of areas, where Δ _morphological measure_ = measure _source area_ – measure _target area_). This resulted in Δ_soma size_, Δ_spine count_, Δ_spine density_, and Δ_tree size_. Each of these difference measures was converted to an undirected variable by computing its absolute value, |Δ_soma size_|, |Δ_spine count_|, |Δ_spine density_|, and |Δ_tree size_|, where appropriate. To relate the four morphological measures to ranked projections strength, we ranked their absolute difference measures separately per target area, analogous to the ranking described for the absolute neuron density ratio above. That is, smaller absolute difference measures were ranked highest (rank 1), and successively larger absolute difference measures were ranked accordingly (increasing rank number).

### Statistical tests

To test groups of projections for equality in their associated structural measures, we computed two-tailed independent samples t-tests and report the t-statistic t, degrees of freedom df and the associated measure of effect size *r*, where *r* = (t^2^ / (t^2^+df))^1/2^. To assess relations between structural measures, connectivity features and area properties, we computed Pearson’s correlation coefficient r. Since ranked projection strength is not an interval measure, we computed Spearman’s rank-correlation coefficient ρ in this case. All tests were conducted with a pre-assigned two-tailed significance level of α = 0.05.

We performed binary logistic regression analyses using projection existence as the binary dependent variable and different combinations of the relative structural measures as covariates. Specifically, we considered |log-ratio_density_|, geodesic distance, |Δ_soma size_|, |Δ_spine count_|, |Δ_spine density_|, and |Δ_tree size_|. The relative structural measures were converted to z-scores, so that the resulting regression coefficients were standardized. We also included a constant in each model. All covariates were entered into the model simultaneously. We report the standardized regression coefficients, the t-statistic, and its associated p-value. We assessed model classification performance in three different ways. First, we calculated the generalized coefficient of determination, R^2^, adjusted for the number of covariates, which indicates which proportion of the variance in the dependent variable is explained by the covariates. Second, we computed the Youden index *J* (Youden, 1950; Fluss et al., 2005), where *J* = sensitivity + specificity -1. By taking into account both sensitivity (true positive rate) and specificity (true negative rate), the Youden index is a comprehensive summary measure of classification performance. It measures how well a binary classifier operates above chance level, where *J* below 0.1 indicates chance performance and *J* = 1 indicates perfect classification. We considered values of *J* below 0.25 to indicate negligible classification performance, values of 0.25 and above to indicate weak performance, values of 0.40 and above to indicate moderate performance, and values of 0.50 and above to indicate good classification performance. Third, we calculated classification accuracy, that is, which proportion of predictions was correct.

## Results

### Macroscopic and microscopic morphological measures are interrelated

The macroscopic and microscopic morphological measures were indeed strongly correlated with each other, with the possible exception of dendritic spine density. When the standard significance threshold of α = 0.05 was applied, all correlations but one (dendritic spine density with dendritic tree size) were statistically significant. Using a Bonferroni correction for multiple tests resulted in an adjusted significance threshold of α_adj_ = 0.05/10 = 0.005. Under this criterion, the significance of all but two correlations remained unaffected. Only the correlation of dendritic spine density with neuron density and with soma size lost statistical significance. Table 1 summarizes these results. Since the structural measures presented such strong interrelations, we chose our methods of analysis accordingly in the following assessment of connectivity features, relying on procedures that took all measures into account conjointly.

**Table 1.** Correlation between structural measures. Pearson correlation coefficients and associated p-values for correlations between neuron density and the morphological measures. Bonferroni correction for multiple tests results in an adjusted significance threshold of α_adj_ = 0.05/10 = 0.005.

|  | Neuron density |  | Soma size |  | Spine count |  | Spine density |  |
| --- | --- | --- | --- | --- | --- | --- | --- | --- |
|  | r | p-value | r | p-value | r | p-value | r | p-value |
| Soma size | -0.63 | 0.0005 | - | - |  |  |  |  |
| Spine count | -0.54 | 0.0028 | 0.56 | 0.0016 | - | - |  |  |
| Spine density | -0.51 | 0.0056 | 0.46 | 0.0130 | 0.91 | <.0001 | - | - |
| Dendritic tree size | -0.57 | 0.0012 | 0.55 | 0.0017 | 0.57 | 0.0005 | 0.32 | 0.0679 |

### Neuron density is most consistently related to the existence of projections

To assess whether the six structural measures were distributed differently across absent and present projections, we computed independent samples t-tests, using the undirected, absolute values of the structural measures. These showed that connected cortical areas had smaller neuron density ratios than areas that were not linked. Similarly, linked areas were separated by smaller distances than unconnected areas. The differences between areas in the four morphological measures were also smaller if areas were connected than if no connection had been found. Of all the tested structural features, effect size was largest for neuron density ratio. These results are summarized in Figure 2 and Table 2.

**Figure 2.**
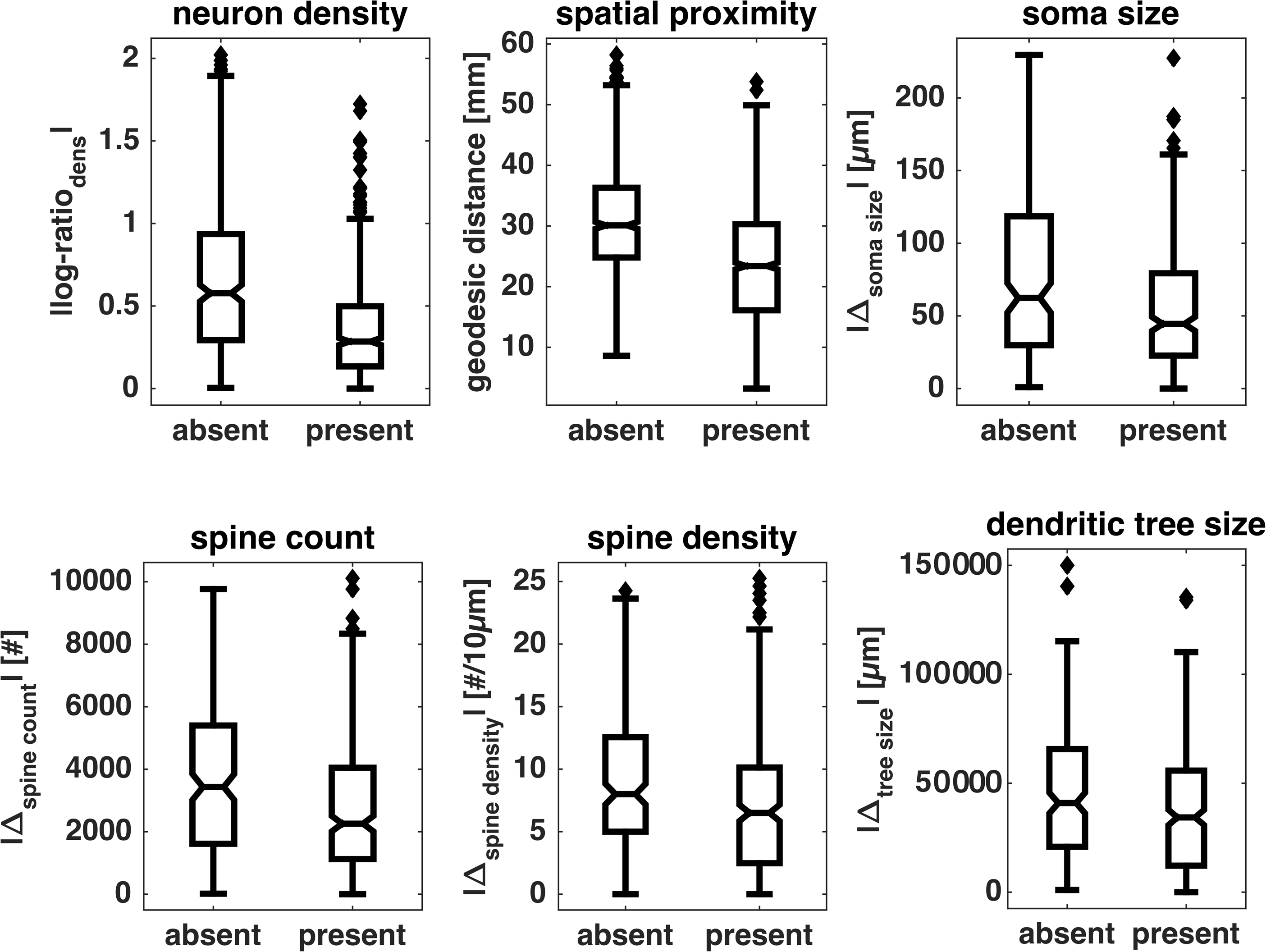
Relative structural measures differ between connected and unconnected pairs of areas. Box plots show distributions of absolute values of relative structural measures for area pairs without (absent) and with (present) a linking projection. Indicated are median (line), interquartile range (box), data range (whiskers) and outliers (diamonds, outside of 2.7 standard deviations). See Table 2 for a summary.

**Table 2.**
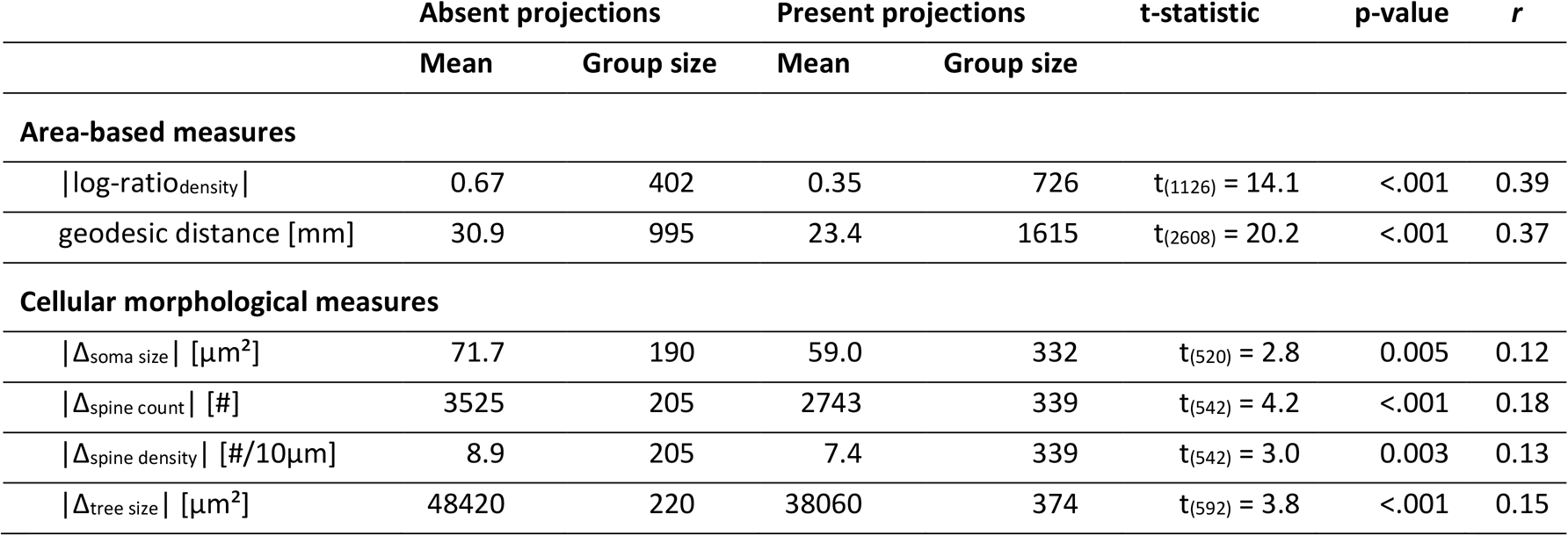
Structural measures in connected and unconnected pairs of areas. Absolute values of relative structural measures are averaged across area pairs without (absent) and with (present) a linking projection. T-statistics, p-values and effect size *r* are results of two-tailed independent samples t-tests comparing the two respective distributions for equal means. See Figure 2 for box plots of the underlying distributions.

|  | Absent projections |  | Present projections |  | t-statistic | p-value | r |
| --- | --- | --- | --- | --- | --- | --- | --- |
|  | Mean | Group size | Mean | Group size |  |  |  |
| Area-based measures |  |  |  |  |  |  |  |
| \|log-ratio<sub>density</sub>\| | 0.67 | 402 | 0.35 | 726 | t<sub>(1126)</sub> = 14.1 | <.001 | 0.39 |
| geodesic distance [mm] | 30.9 | 995 | 23.4 | 1615 | t<sub>(2608)</sub> = 20.2 | <.001 | 0.37 |
| Cellular morphological measures |  |  |  |  |  |  |  |
| \|Δsoma size\| [μm<sup>2</sup>] | 71.7 | 190 | 59.0 | 332 | t<sub>(520)</sub> = 2.8 | 0.005 | 0.12 |
| \|Δspine count\| [#] | 3525 | 205 | 2743 | 339 | t<sub>(542)</sub> = 4.2 | <.001 | 0.18 |
| \|Δspine density\| [#/<sup>10</sup>μm] | 8.9 | 205 | 7.4 | 339 | t<sub>(542)</sub> = 3.0 | 0.003 | 0.13 |
| \|Δtree size\| [μm<sup>2</sup>] | 48420 | 220 | 38060 | 374 | t<sub>(592)</sub> = 3.8 | <.001 | 0.15 |

We used binary logistic regression to probe how well the structural measures could account for the existence of projections (Figure 3, Table 3). First, we tested each structural measure individually (logistic regressions 1-6), computing logistic regressions with either absolute neuron density ratio, geodesic distance, absolute soma size difference, absolute spine count difference, absolute spine density difference, or absolute dendritic tree size difference as the only covariate (plus an intercept term). We also computed a null model that contained only the intercept term (logistic regression 7) and represented chance performance. Second, we included all six measures in conjunction as covariates (logistic regression 8). Third, we removed each measure separately from the conjoint set of covariates (logistic regressions 9-14), so that we computed six logistic regressions with five of the six structural measures as covariates. We assessed model classification performance through the adjusted generalized R^2^, the Youden index *J*, and prediction accuracy. As presented in Table 3, each measure contributed significantly (as indicated by the p-value) to the model performance if it was the only covariate. However, all three performance measures (R^2^, *J* and accuracy) indicated that classification performance was essentially at chance level for all four cellular morphological measures. Both the neuron density ratio and geodesic distance reached weak classification performance on their own, as indicated by R^2^ and *J*. Accuracy was slightly above chance for geodesic distance and clearly above chance level for the neuron density ratio.

**Table 3.**
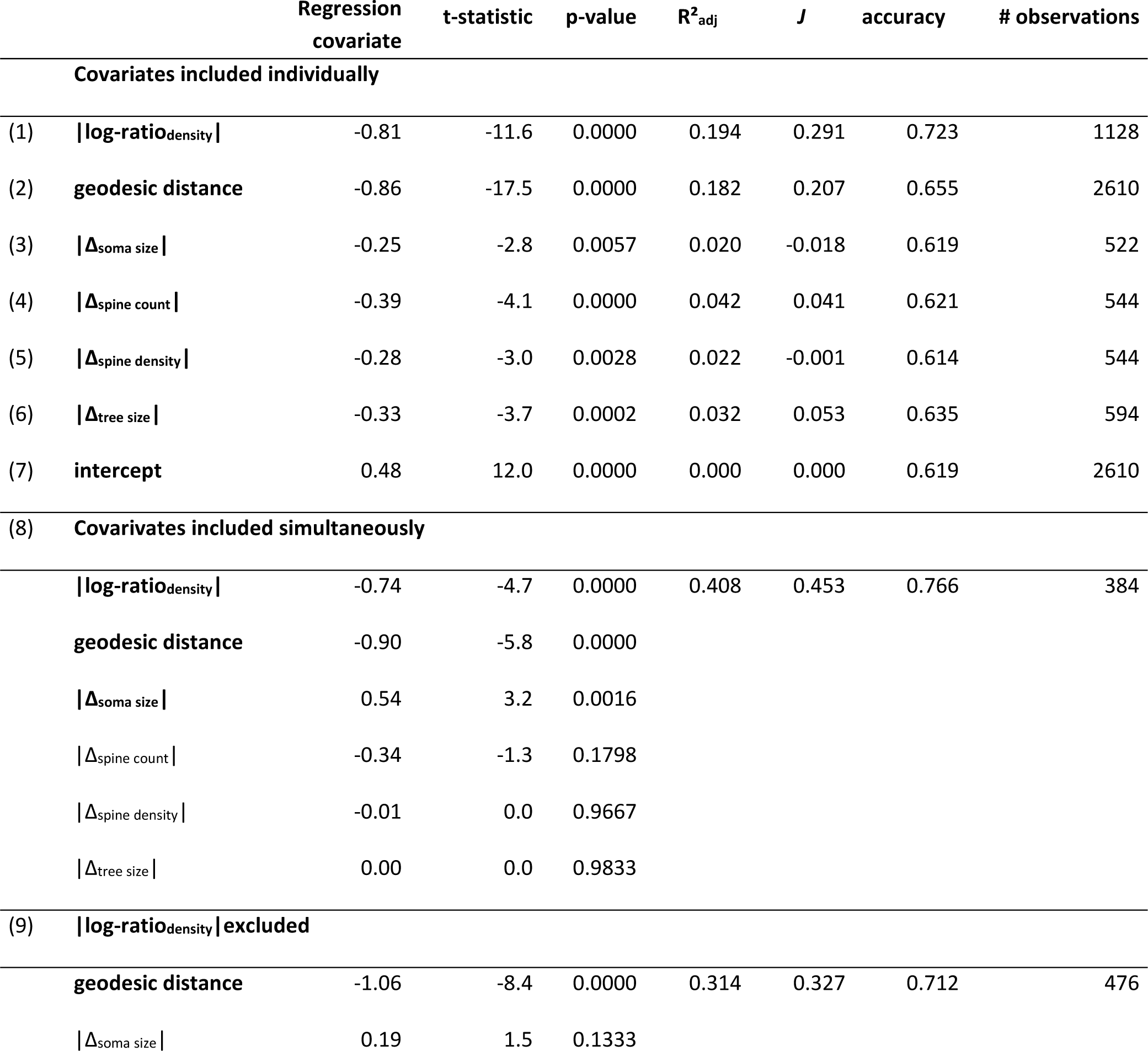
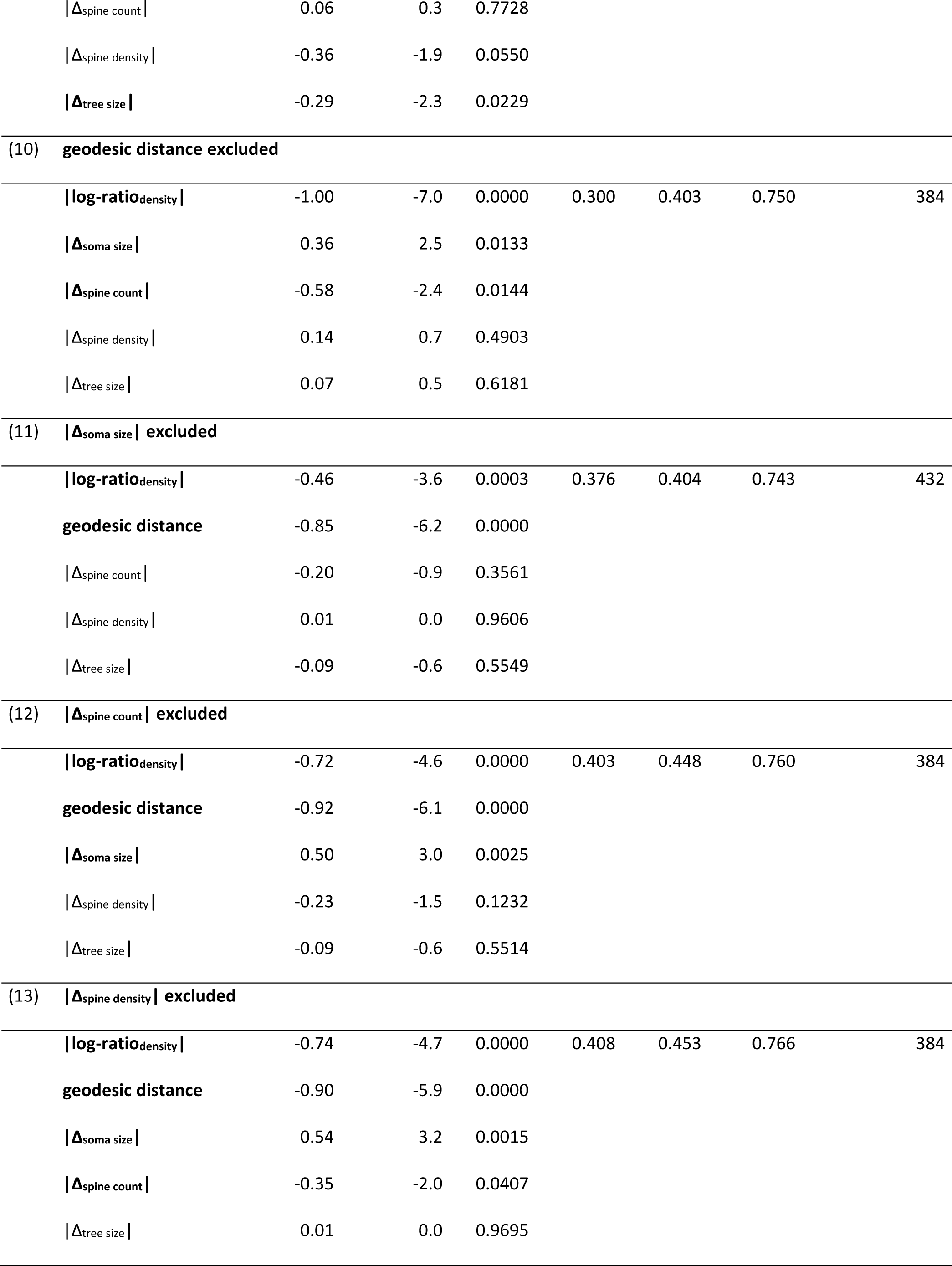
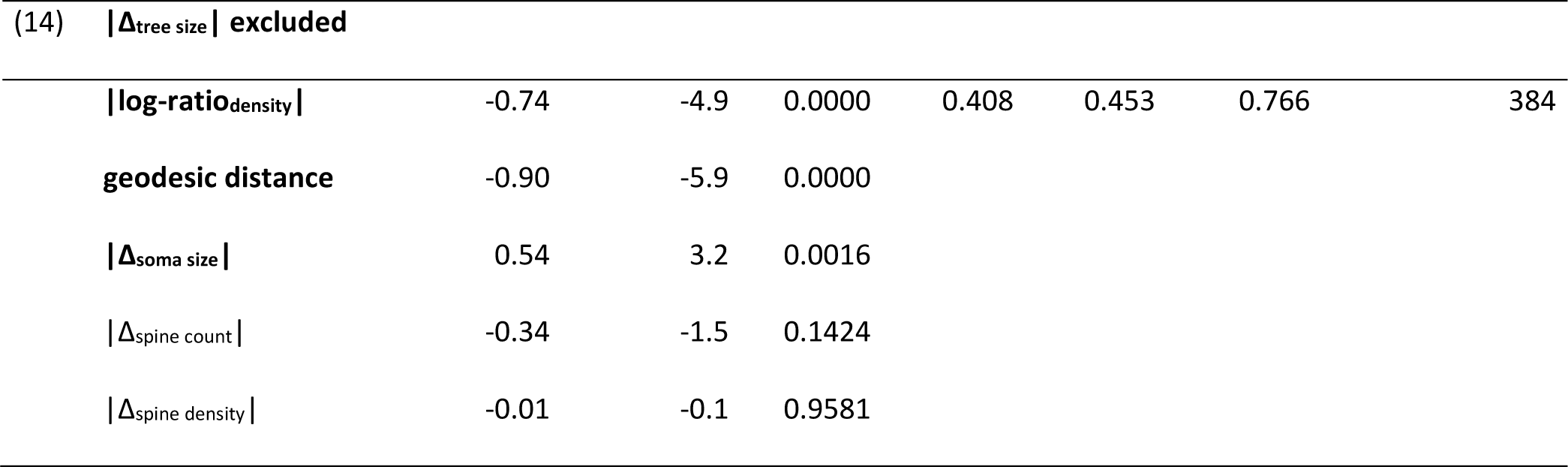
Classification of connection existence by logistic regression. We performed binary logistic regression analyses (enumerated in brackets), each including a different set of the structural measures as covariates and connectivity (grouped into ‘absent’ and ‘present’ connections) as the dependent variable. Bold-faced covariates significantly contributed to classification performance as indicated by the p-value. Across all regression analyses, absolute neuron density ratio, geodesic distance and soma size difference emerged as meaningful predictors.

|  |  | Regression covariate | t-statistic | p-value | R<sup>2</sup><sub>adj</sub> | J | accuracy | # observations |
| --- | --- | --- | --- | --- | --- | --- | --- | --- |
| **Covariates included individually** |  |  |  |  |  |  |  |  |
| (1) | **\|log-ratio<sub>density</sub>\|** | -0.81 | -11.6 | 0.0000 | 0.194 | 0.291 | 0.723 | 1128 |
| (2) | **geodesic distance** | -0.86 | -17.5 | 0.0000 | 0.182 | 0.207 | 0.655 | 2610 |
| (3) | **\|Δ<sub>soma size</sub>\|** | -0.25 | -2.8 | 0.0057 | 0.020 | -0.018 | 0.619 | 522 |
| (4) | **\|Δ<sub>spine count</sub>\|** | -0.39 | -4.1 | 0.0000 | 0.042 | 0.041 | 0.621 | 544 |
| (5) | **\|Δ<sub>spine density</sub>\|** | -0.28 | -3.0 | 0.0028 | 0.022 | -0.001 | 0.614 | 544 |
| (6) | **\|Δ<sub>tree size</sub>\|** | -0.33 | -3.7 | 0.0002 | 0.032 | 0.053 | 0.635 | 594 |
| (7) | **intercept** | 0.48 | 12.0 | 0.0000 | 0.000 | 0.000 | 0.619 | 2610 |
| **(8) Covarivates included simultaneously** |  |  |  |  |  |  |  |  |
|  | **\|log-ratio<sub>density</sub>\|** | -0.74 | -4.7 | 0.0000 | 0.408 | 0.453 | 0.766 | 384 |
|  | **geodesic distance** | -0.90 | -5.8 | 0.0000 |  |  |  |  |
|  | **\|Δ<sub>soma size</sub>\|** | 0.54 | 3.2 | 0.0016 |  |  |  |  |
|  | **\|Δ<sub>spine count</sub>\|** | -0.34 | -1.3 | 0.1798 |  |  |  |  |
|  | **\|Δ<sub>spine density</sub>\|** | -0.01 | 0.0 | 0.9667 |  |  |  |  |
|  | **\|Δ<sub>tree size</sub>\|** | 0.00 | 0.0 | 0.9833 |  |  |  |  |
| **(9) \|log-ratio<sub>density</sub>\| excluded** |  |  |  |  |  |  |  |  |
|  | **geodesic distance** | -1.06 | -8.4 | 0.0000 | 0.314 | 0.327 | 0.712 | 476 |
|  | **\|Δ<sub>soma size</sub>\|** | 0.19 | 1.5 | 0.1333 |  |  |  |  |

|  |  |  |  |  |  |  |  |
| --- | --- | --- | --- | --- | --- | --- | --- |
| **\|Δspine count\|** | 0.06 | 0.3 | 0.7728 |  |  |  |  |
| **\|Δspine density\|** | -0.36 | -1.9 | 0.0550 |  |  |  |  |
| **\|Δtree size\|** | -0.29 | -2.3 | 0.0229 |  |  |  |  |
| **(10) geodesic distance excluded** |  |  |  |  |  |  |  |
| **\|log-ratio<sub>density</sub>\|** | -1.00 | -7.0 | 0.0000 | 0.300 | 0.403 | 0.750 | 384 |
| **\|Δsoma size\|** | 0.36 | 2.5 | 0.0133 |  |  |  |  |
| **\|Δspine count\|** | -0.58 | -2.4 | 0.0144 |  |  |  |  |
| **\|Δspine density\|** | 0.14 | 0.7 | 0.4903 |  |  |  |  |
| **\|Δtree size\|** | 0.07 | 0.5 | 0.6181 |  |  |  |  |
| **(11) \|Δsoma size\| excluded** |  |  |  |  |  |  |  |
| **\|log-ratio<sub>density</sub>\|** | -0.46 | -3.6 | 0.0003 | 0.376 | 0.404 | 0.743 | 432 |
| **geodesic distance** | -0.85 | -6.2 | 0.0000 |  |  |  |  |
| **\|Δspine count\|** | -0.20 | -0.9 | 0.3561 |  |  |  |  |
| **\|Δspine density\|** | 0.01 | 0.0 | 0.9606 |  |  |  |  |
| **\|Δtree size\|** | -0.09 | -0.6 | 0.5549 |  |  |  |  |
| **(12) \|Δspine count\| excluded** |  |  |  |  |  |  |  |
| **\|log-ratio<sub>density</sub>\|** | -0.72 | -4.6 | 0.0000 | 0.403 | 0.448 | 0.760 | 384 |
| **geodesic distance** | -0.92 | -6.1 | 0.0000 |  |  |  |  |
| **\|Δsoma size\|** | 0.50 | 3.0 | 0.0025 |  |  |  |  |
| **\|Δspine density\|** | -0.23 | -1.5 | 0.1232 |  |  |  |  |
| **\|Δtree size\|** | -0.09 | -0.6 | 0.5514 |  |  |  |  |
| **(13) \|Δspine density\| excluded** |  |  |  |  |  |  |  |
| **\|log-ratio<sub>density</sub>\|** | -0.74 | -4.7 | 0.0000 | 0.408 | 0.453 | 0.766 | 384 |
| **geodesic distance** | -0.90 | -5.9 | 0.0000 |  |  |  |  |
| **\|Δsoma size\|** | 0.54 | 3.2 | 0.0015 |  |  |  |  |
| **\|Δspine count\|** | -0.35 | -2.0 | 0.0407 |  |  |  |  |
| **\|Δtree size\|** | 0.01 | 0.0 | 0.9695 |  |  |  |  |

| (14) $\vert \Delta_{\text{tree size}}\vert $ excluded | | | | | | | |
| --- | --- | --- | --- | --- | --- | --- | --- |
| $\vert \log\text{-ratio}_{\text{density}}\vert $ | -0.74 | -4.9 | 0.0000 | 0.408 | 0.453 | 0.766 | 384 |
| geodesic distance | -0.90 | -5.9 | 0.0000 |  |  |  |  |
| $\vert \Delta_{\text{soma size}}\vert $ | 0.54 | 3.2 | 0.0016 | | | | |
| $\vert \Delta_{\text{spine count}}\vert $ | -0.34 | -1.5 | 0.1424 | | | | |
| $\vert \Delta_{\text{spine density}}\vert $ | -0.01 | -0.1 | 0.9581 | | | | |

Furthermore, if all measures were included as covariates simultaneously, both macroscopic area-based measures contributed significantly to the model performance, while the only cellular morphological measure that remained significant was soma size difference. Model performance increased to moderate levels as indicated by all three performance measures. From the overall analysis of model performance, the neuron density ratio emerged as the most important predictive factor for projection existence. This was indicated by the fact that the decline in model performance was largest for the exclusion of neuron density ratio (9), compared to the exclusion of the other five measures (10-14). Logistic regressions 10 and 11 demonstrated that geodesic distance and soma size difference also added meaningful information regarding the existence of projections. The other three cellular morphological measures did not contribute any additional information, as can be seen in logistic regressions 12 to 14, where model performance was essentially identical to the full model, although either spine count difference, spine density difference or dendritic tree size difference were excluded. These observations support our conclusion that neuron density was the neural structural measure that principally related to the existence of connections.

### Distance and dendritic tree size are related to projection strength

The strength of cortical projections has been shown to diminish with greater distance of the connected areas (e.g., Ercsey-Ravasz et al., 2013). In the next step, we related all area-based macroscopic as well as cell-based microscopic measures to the strength of projections, ln(FLNe), using the undirected, absolute values of the structural measures. Figure 4 and Table 4 show that each measure individually was weakly to moderately correlated with projection strength, with the exception of soma size difference. The strongest correlation was found for geodesic distance. For the absolute neuron density ratio, the negative correlation coefficient signified that areas that were similar in their neuron density (i.e., with a small absolute density ratio) were linked by stronger projections than areas that were less similar in their density (i.e., with a large absolute density ratio). Since most measures were correlated with each other, we entered all measures into a partial Pearson correlation to assess their relative contribution to projection strength if the other measures were controlled for. As shown in Table 4, when all other measures were controlled for, geodesic distance and dendritic tree size difference retained their correlation to projection strength, with the respective correlation coefficients even gaining slightly in magnitude. The third measure that was significantly correlated with projection strength in the partial correlation was soma size difference. Neuron density ratio and spine count difference clearly did not correlate with projection strength if all measures were considered simultaneously. The correlation of spine density difference was close to remaining significant; the magnitude, however, was very weak. These results were mirrored in our analysis of ranked projection strength (Figure 4, Table 4), where we ranked both the strength of projections (i.e., incoming projections were ordered from strongest to weakest) and the difference measures per target area. Again, all measures were significantly correlated with projection strength if considered individually. The strongest association was again observed for geodesic distance. However, if all other measures were accounted for in a partial correlation, only geodesic distance and dendritic tree size difference remained significant. All significant correlation coefficients were positive, indicating that weaker projections (with higher rank numbers) were associated with larger absolute differences in the structural measures (with higher rank numbers) between connected areas.

**Figure 4.**
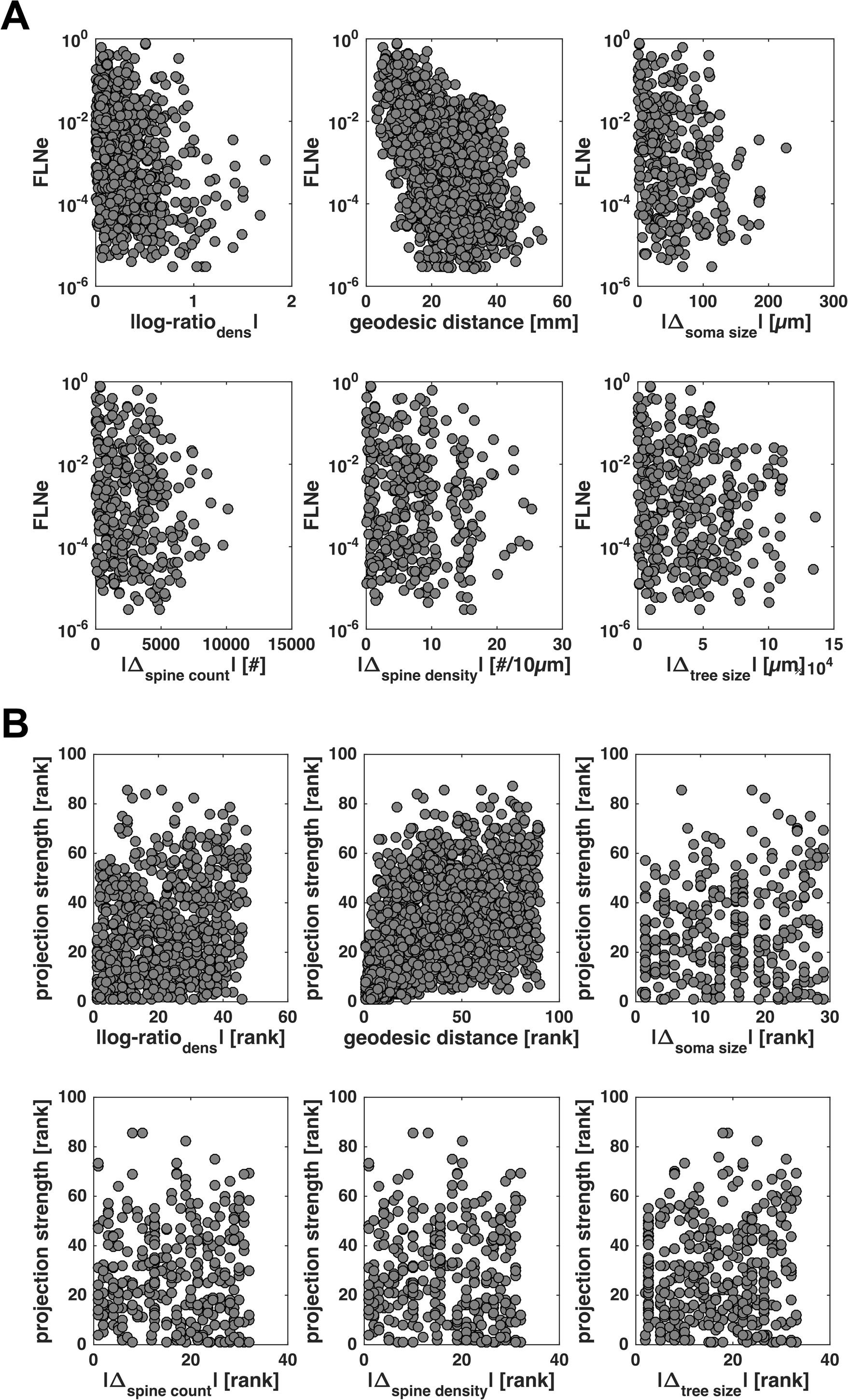
Projection strength varies with relative structural measures. (A) Projection strength for individual projections, FLNe, is shown across absolute values of relative structural measures. (B) Ranked projection strength of individual projections is shown across ranked absolute values of relative structural measures. See Table 4 for correlation statistics.

**Table 4.** Correlation between projection strength and structural measures. Pearson correlation coefficients and associated p-values for correlations between projection strength, expressed either as ln(FLNe) or as ranked strengths, and absolute values of relative structural measures or ranked absolute values of relative structural measures. Correlations were assessed both for each measure independently (individual correlation) and while accounting for all other five measures (partial correlation). See Figure 4 for scatter plots of the underlying distributions.

| **FLNe** | **Individual correlation** |  | **Partial correlation** |  |
| --- | --- | --- | --- | --- |
|  | r | p-value | r | p-value |
| \|log-ratio<sub>density</sub>\| | -0.25 | <0.0001 | -0.08 | 0.2158 |
| geodesic distance | -0.49 | <0.0001 | -0.53 | <0.0001 |
| \|$\Delta_{\text{soma size}}$\| | -0.10 | 0.0787 | 0.14 | 0.0319 |
| \|$\Delta_{\text{spine count}}$\| | -0.17 | 0.0014 | 0.00 | 0.9716 |
| \|$\Delta_{\text{spine density}}$\| | -0.12 | 0.0332 | -0.13 | 0.0527 |
| \|$\Delta_{\text{tree size}}$\| | -0.13 | 0.0094 | -0.19 | 0.0028 |
| **ranked strength** | **$\rho$** | **p-value** | **$\rho$** | **p-value** |
| ranked \|log-ratio<sub>density</sub>\| | 0.24 | <0.0001 | 0.11 | 0.0800 |
| ranked geodesic distance | 0.51 | <0.0001 | 0.52 | <0.0001 |
| ranked \|$\Delta_{\text{soma size}}$\| | 0.17 | 0.0019 | -0.12 | 0.0656 |
| ranked \|$\Delta_{\text{spine count}}$\| | 0.15 | 0.0050 | -0.01 | 0.8921 |
| ranked \|$\Delta_{\text{spine density}}$\| | 0.12 | 0.0256 | 0.13 | 0.0533 |
| ranked \|$\Delta_{\text{tree size}}$\| | 0.17 | 0.0013 | 0.18 | 0.0047 |

### Neuron density is consistently related to laminar patterns of projection origins

Our analyses of the extensive projection origin data provided in Chaudhuri et al. (2015) show that neuron density is consistently related to the laminar pattern of corticocortical projection origins. As seen from Figure 5 and Table 5, all directed measures were weakly to moderately correlated with the laminar pattern of projections expressed as *N*_SG_%. For the neuron density ratio, the positive correlation coefficient indicated that projections from less dense areas to denser areas had a more infragranular origin, while projections from denser to less dense areas had predominantly supragranular origins. For the cellular morphological measures, the negative correlation coefficient indicated the reverse relationship. However, in a partial Pearson correlation of *N*_SG_% with all measures, except geodesic distance, only neuron density ratio and spine count difference retained significance. Although the correlation with soma size difference was below the significance threshold, the correlation coefficient changed its sign while remaining at a weak magnitude, indicating that the correlation was volatile and not reliable. Geodesic distance, an undirected measure, was tested for a correlation with an indicator of deviation from bilaminar projection patterns, |*N*_SG_%|. This correlation only reached a weak magnitude. To test the effect of additionally controlling the partial Pearson correlation for geodesic distance, we computed a partial Pearson correlation of |*N*_SG_%| with geodesic distance as well as the absolute values of the other five measures. Here, absolute neuron density ratio was the only measure that remained significant, retaining its moderate magnitude. In conclusion, the variable found to be significantly and most strongly associated with the laminar pattern of projections across all variations of correlating the measures was the neuron density ratio.

**Figure 5.**
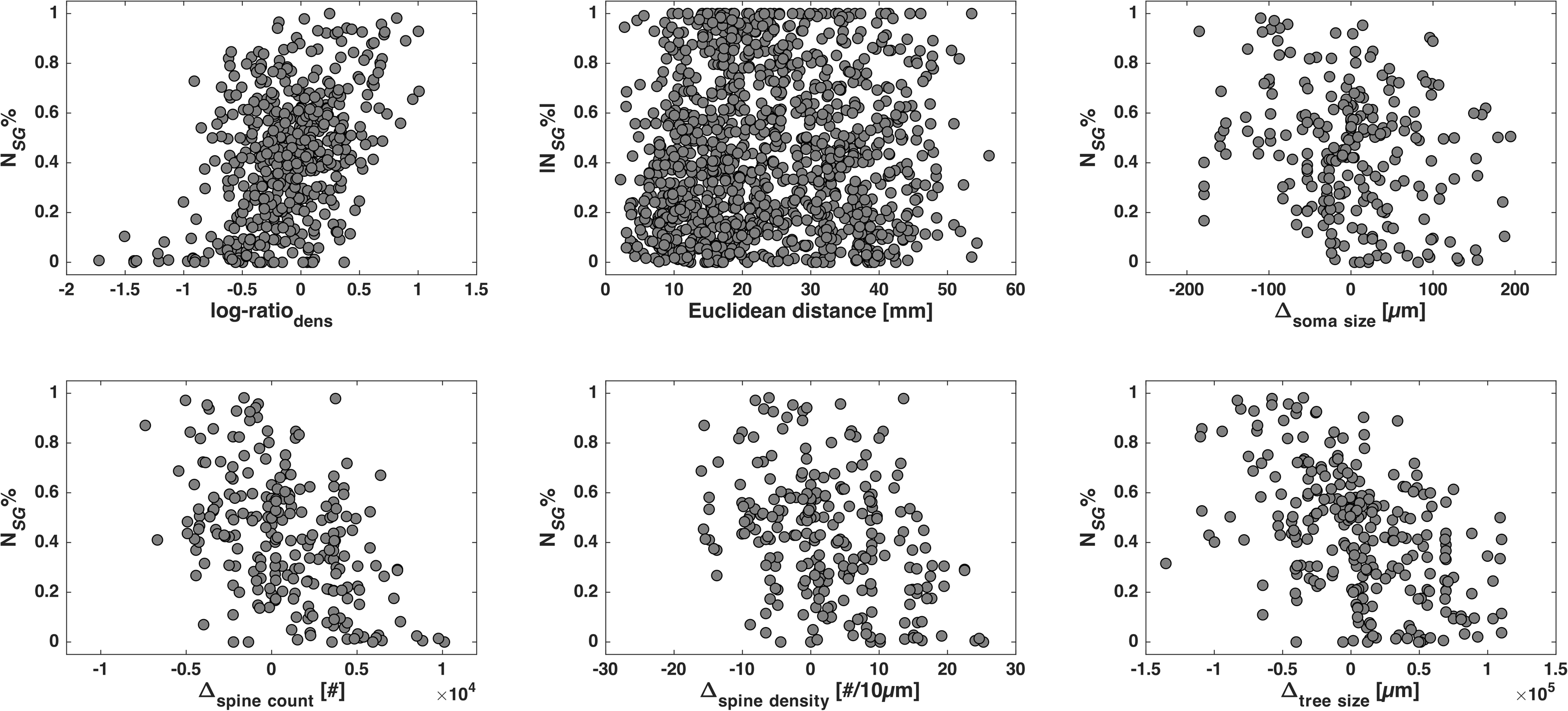
Laminar projection patterns vary with relative structural measures. Fraction of supragranularly labelled neurons for individual projections, *N*_SG_%, is shown across relative structural measures. Note that for geodesic distance, the measure of deviation from columnar laminar patterns |*N*_SG_%| is shown instead. See Table 5 for correlation statistics.

**Table 5.** Correlation between laminar projection patterns and structural measures. Pearson correlation coefficients and associated p-values for correlations between *N*_SG_% and relative structural measures. Correlations were assessed both for each measure independently (individual correlation with *N*_SG_%) and while accounting for four other measures (partial correlation with *N*_SG_%). Geodesic distance could not be correlated with *N*_SG_%, because it is an undirected measure. Instead, the individual correlation of geodesic distance with laminar patterns (marked ‘*’) was computed using |*N*_SG_%|, which indicates deviation from bilaminar projection patterns. Accordingly, we also computed a partial correlation accounting for all six structural measures at once, which had to include the absolute values of the relative structural measures (partial correlation with |*N*_SG_%|). See Figure 5 for scatter plots of the underlying distributions.

| | Individual correlation<br>with $N_{SG}\%$ | | Partial correlation<br>with $N_{SG}\%$ | | | Partial correlation<br>with $\vert N_{SG}\%\vert $ | |
| --- | --- | --- | --- | --- | --- | --- | --- |
|  | r | p-value | r | p-value |  | r | p-value |
| log-ratio<sub>density</sub> | 0.43 | <.0001 | 0.40 | <.0001 | $\vert \text{log-ratio}_{\text{density}}\vert $ | 0.38 | <.0001 |
| geodesic distance* | 0.12 | 0.0001 | -/- | -/- | geodesic distance | -0.07 | 0.3651 |
| $\Delta_{\text{soma size}}$ | -0.21 | 0.0009 | 0.20 | 0.0090 | $\vert \Delta_{\text{soma size}}\vert $ | -0.03 | 0.7406 |
| $\Delta_{\text{spine count}}$ | -0.47 | <.0001 | -0.18 | 0.0206 | $\vert \Delta_{\text{spine count}}\vert $ | 0.09 | 0.2708 |
| $\Delta_{\text{spine density}}$ | -0.35 | <.0001 | 0.08 | 0.2910 | $\vert \Delta_{\text{spine density}}\vert $ | 0.07 | 0.3658 |
| $\Delta_{\text{tree size}}$ | -0.47 | <.0001 | -0.11 | 0.1428 | $\vert \Delta_{\text{tree size}}\vert $ | 0.02 | 0.7975 |

### Neuron density is correlated with the number of connections of cortical areas

As in previous reports (Scholtens et al., 2014; Beul et al., 2017), we examined the correlation of structural measures with overall area degree, the number of connections a cortical region maintains (within the 29×29 edge-complete subgraph). We tested five of the six measures. Spatial proximity was excluded, because geodesic distance is inherently relational and cannot be related to a measure pertaining to a single area. Figure 6 and Table 6 show that of the five tested measures, only neuron density was significantly correlated with overall area degree (this correlation of neuron density was reported previously in Beul et al., 2017), although the correlation of dendritic tree size with overall degree was very close to significant. Using a Bonferroni correction for multiple tests resulted in an adjusted significance threshold of α_adj_ = 0.05/5 = 0.01. Neuron density remained significantly correlated with overall area degree also under this criterion. Similarly, in a partial Pearson correlation, only neuron density was significantly correlated with overall area degree, at a strong magnitude.

**Figure 6.**
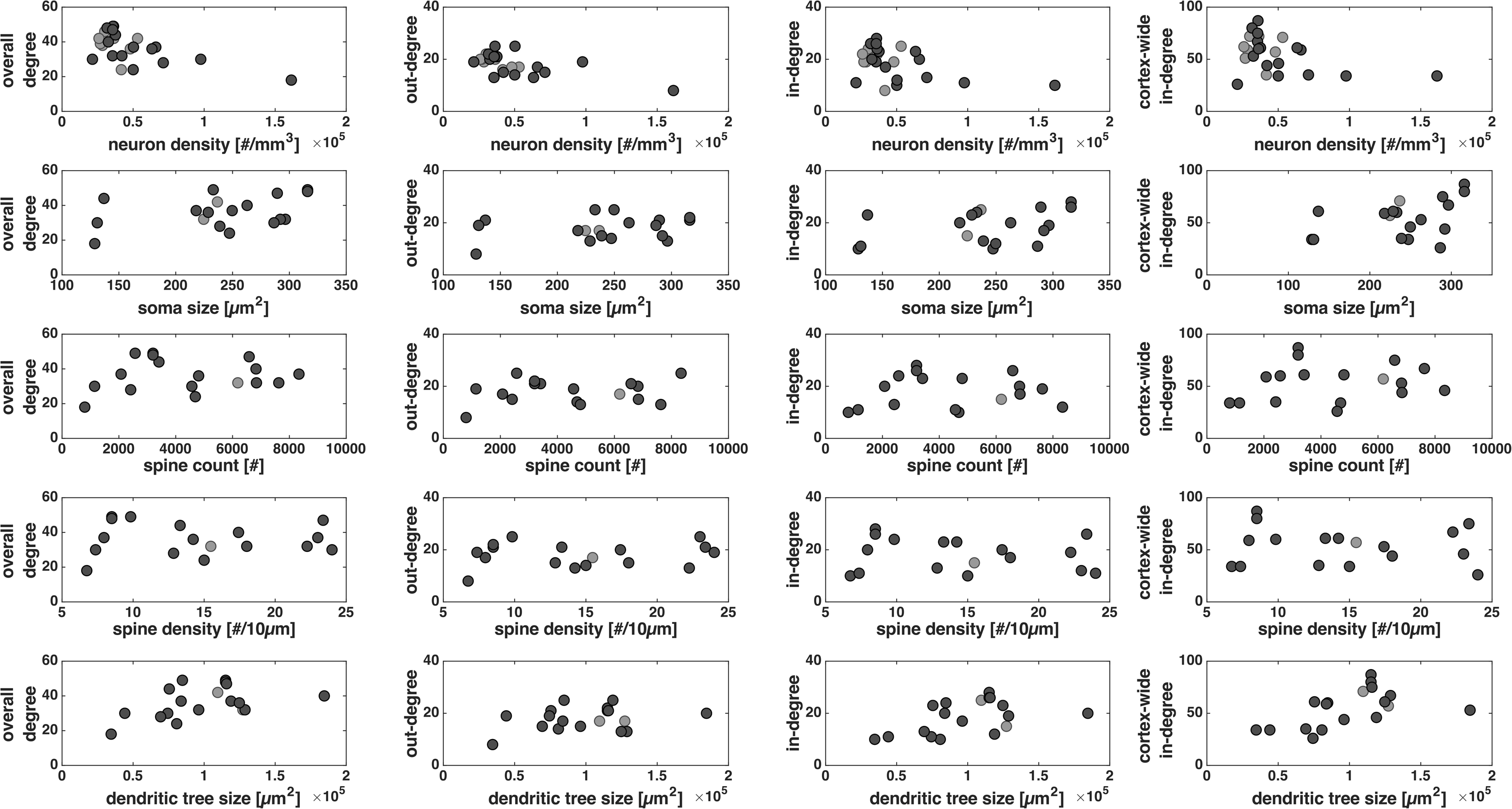
Area degree varies with structural measures. The number of projections an area maintains, area degree, is shown across structural measures for each area. Depicted are overall degree, out-degree and in-degree on the 29×29 edge-complete subgraph, as well as cortex-wide in-degree. See Table 6 for correlation statistics. All data points (black and grey) were considered in the individual correlations reported there, while only black data points were considered in the partial correlation.

**Table 6.**
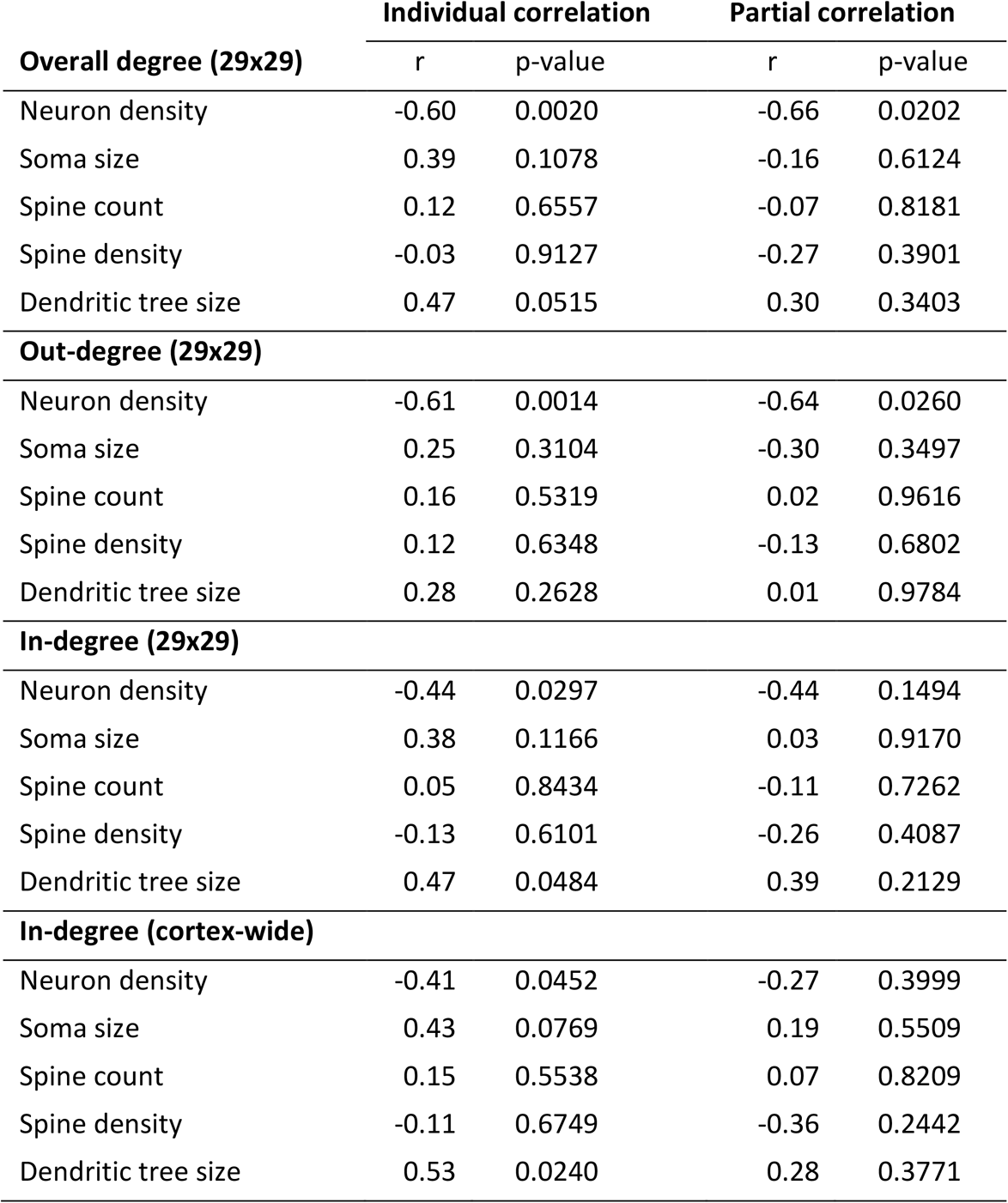
Correlation between area degree and structural measures. Pearson correlation coefficients and associated p-values for correlations between the structural measures for each area and overall area degree (total number of maintained connections), out-degree, in-degree or cortex-wide in-degree. Correlations were assessed both for each measure independently (individual correlation) and while accounting for the other five measures (partial correlation). Geodesic distance could not be included because it is a relational property which is not defined for individual areas. Bonferroni correction for multiple tests results in an adjusted significance threshold of α_adj_ = 0.05/5 = 0.01 for the individual correlations. The individual correlation of neuron density with degree has been reported previously (Beul et al., 2017). See Figure 6 for scatter plots of the underlying distributions.

|  | Individual correlation |  | Partial correlation |  |
| --- | --- | --- | --- | --- |
|  | r | p-value | r | p-value |
| **Overall degree (29x29)** |  |  |  |  |
| Neuron density | -0.60 | 0.0020 | -0.66 | 0.0202 |
| Soma size | 0.39 | 0.1078 | -0.16 | 0.6124 |
| Spine count | 0.12 | 0.6557 | -0.07 | 0.8181 |
| Spine density | -0.03 | 0.9127 | -0.27 | 0.3901 |
| Dendritic tree size | 0.47 | 0.0515 | 0.30 | 0.3403 |
| **Out-degree (29x29)** |  |  |  |  |
| Neuron density | -0.61 | 0.0014 | -0.64 | 0.0260 |
| Soma size | 0.25 | 0.3104 | -0.30 | 0.3497 |
| Spine count | 0.16 | 0.5319 | 0.02 | 0.9616 |
| Spine density | 0.12 | 0.6348 | -0.13 | 0.6802 |
| Dendritic tree size | 0.28 | 0.2628 | 0.01 | 0.9784 |
| **In-degree (29x29)** |  |  |  |  |
| Neuron density | -0.44 | 0.0297 | -0.44 | 0.1494 |
| Soma size | 0.38 | 0.1166 | 0.03 | 0.9170 |
| Spine count | 0.05 | 0.8434 | -0.11 | 0.7262 |
| Spine density | -0.13 | 0.6101 | -0.26 | 0.4087 |
| Dendritic tree size | 0.47 | 0.0484 | 0.39 | 0.2129 |
| **In-degree (cortex-wide)** |  |  |  |  |
| Neuron density | -0.41 | 0.0452 | -0.27 | 0.3999 |
| Soma size | 0.43 | 0.0769 | 0.19 | 0.5509 |
| Spine count | 0.15 | 0.5538 | 0.07 | 0.8209 |
| Spine density | -0.11 | 0.6749 | -0.36 | 0.2442 |
| Dendritic tree size | 0.53 | 0.0240 | 0.28 | 0.3771 |

In addition to overall area degree, we also considered incoming and outgoing connections separately as in-degree and out-degree (Figure 6, Table 6). The only measure that was correlated with out-degree (within the 29×29 edge-complete subgraph), either individually or in a partial correlation, was neuron density, at a strong magnitude, mirroring the results for overall area degree. In-degree (both within the 29×29 edge-complete subgraph and cortex-wide across all 91 areas in the M132 parcellation) was moderately to strongly and significantly correlated with both neuron density and dendritic tree size if the measures were correlated individually. However, in a partial correlation no correlation remained significant. Both neuron density and dendritic tree size retained correlation coefficients of moderate magnitudes, however.

### Discarding very weak projections does not affect the observed relationships

To exclude the possibility that the reported results were mainly driven by very weak projections that are potentially spurious, we repeated all analyses with a smaller connectivity data set, from which projections that did not have at least five constituent axons were excluded (Supplementary Tables 2 to 5). None of the reported results was affected substantially, and the same conclusions can be drawn as from the main analyses.

### Discussion

The extent to which cortical architecture determines the organization of structural connectivity in the cerebral cortex has been examined from a variety of macroscopic and microscopic perspectives (Hilgetag and Grant, 2010; Scholtens et al., 2014; Beul et al., 2015; Hilgetag et al., 2016; Beul et al., 2017; for a review see Barbas, 2015). Here, we explored the relative explanatory power of six structural measures with regard to the organization of corticocortical connections in the macaque cortex. These architectonic measures were examined individually in previous reports (Scholtens et al., 2014; Hilgetag et al., 2016; Beul et al., 2017) and fall into two broad categories: The first group consists of the macroscopic measures of neuron density and spatial proximity. The second group comprises the microscopic cellular morphological measures of soma size, total dendritic spine count, peak dendritic spine density, and dendritic tree size, all measured in L3 cortical pyramidal neurons. We considered these measures in conjunction, to assess how they relate to each other as well as to establish which of them carried the most weight for explaining fundamental organizational aspects of the primate cortical connectome. We found that all morphological measures were strongly correlated with neuron density as well as mostly interrelated among each other (Table 1). Moreover, all six measures diverged depending on whether areas were linked by a projection or not (Figure 2, Table 2). This finding raises the question of whether all of these measures contribute equally to the discrimination of corticocortical connectivity, or whether some of them are redundant, being dependent on other factors, and supply no additional information. Systematic analysis by multivariate logistic regression (Figure 3, Table 3) revealed that of the six measures, only three carried significant information allowing the prediction of projection existence. These were neuron density, geodesic distance and L3 pyramidal cell soma size. The other three cellular morphological measures did not add any information. Of the three significant predictive factors, neuron density emerged as the most relevant, reaching the best classification performance on its own and resulting in the largest decline in classification performance if excluded. This finding suggests that neuron density was the most informative neural measure regarding the existence of projections.

**Figure 3.**
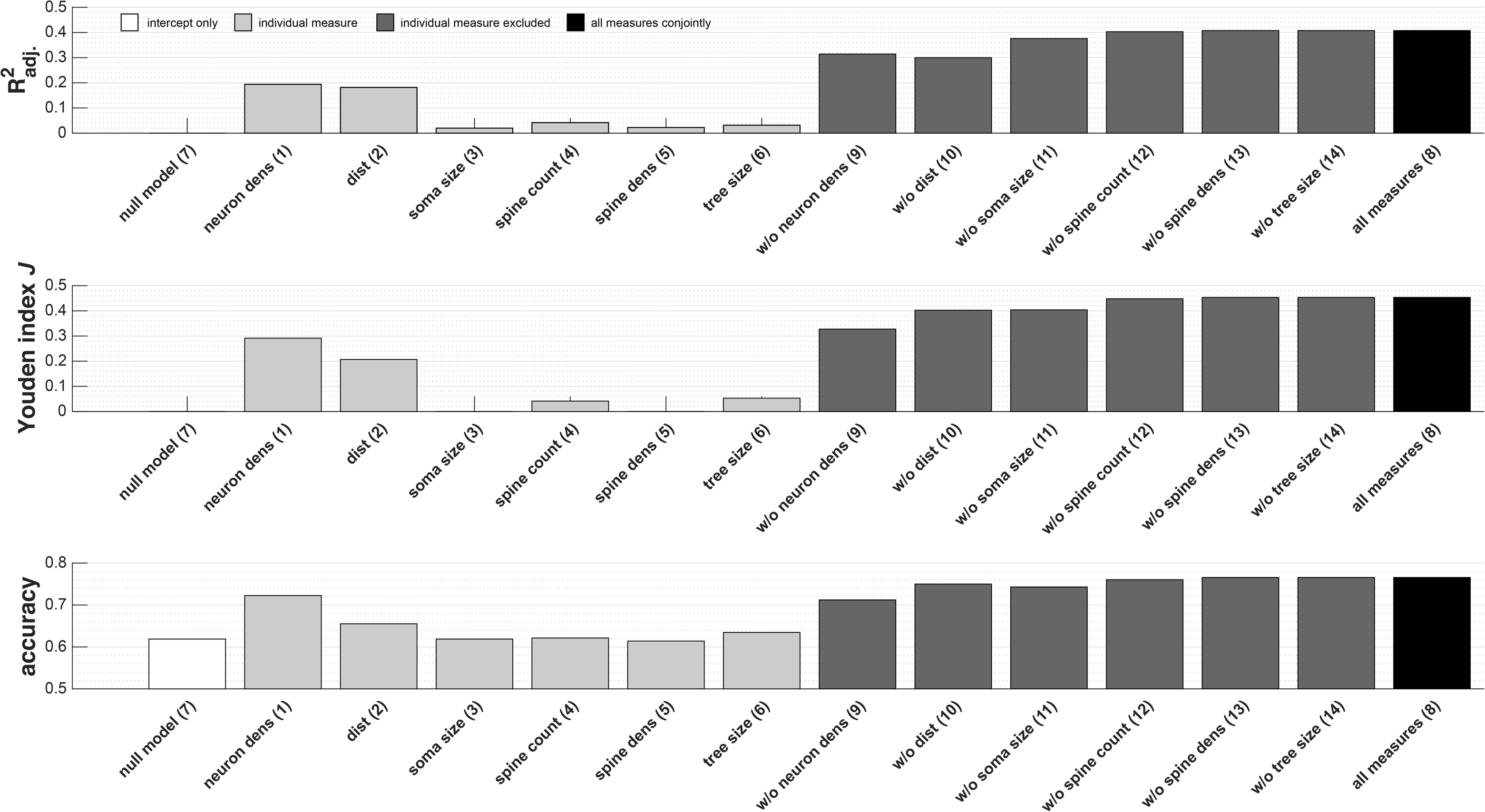
Classification of connection existence by logistic regression. Projection existence was classified using binary logistic regression analyses, which included different combinations of the structural measures as covariates. The classification performance measures adjusted R^2^, Youden index *J* and prediction accuracy are depicted for all logistic regressions. See Table 3 for regression coefficients and test statistics of all regression analyses. Enumeration corresponds to Table 3.

We further found that, even if all other structural measures were controlled for, geodesic distance remained strongly and dendritic tree size difference remained weakly correlated with projection strength (Table 4). This is in line with previous reports showing projection strength to decline as distance between connected areas increases (Ercsey-Ravasz et al., 2013; Markov et al., 2013b).

Additionally, the laminar patterns of projection origins were correlated with the neuron density ratio (Figure 5, Table 5), but not as strongly or consistently with the other five measures. This corroborates previous findings on the importance of architectonic differentiation regarding the laminar distribution of axon terminals (e.g., Barbas, 1986; Beul et al., 2015; Hilgetag et al., 2016). Figure 7 illustrates the observed patterns of projection origins in the context of the five structural measures. Furthermore, the observed lack of a meaningful correlation between laminar projection patterns and spatial proximity directly contradicts the hypothesis that physical distance has a crucial role in affecting laminar patterns (Salin and Bullier, 1995).

**Figure 7.**
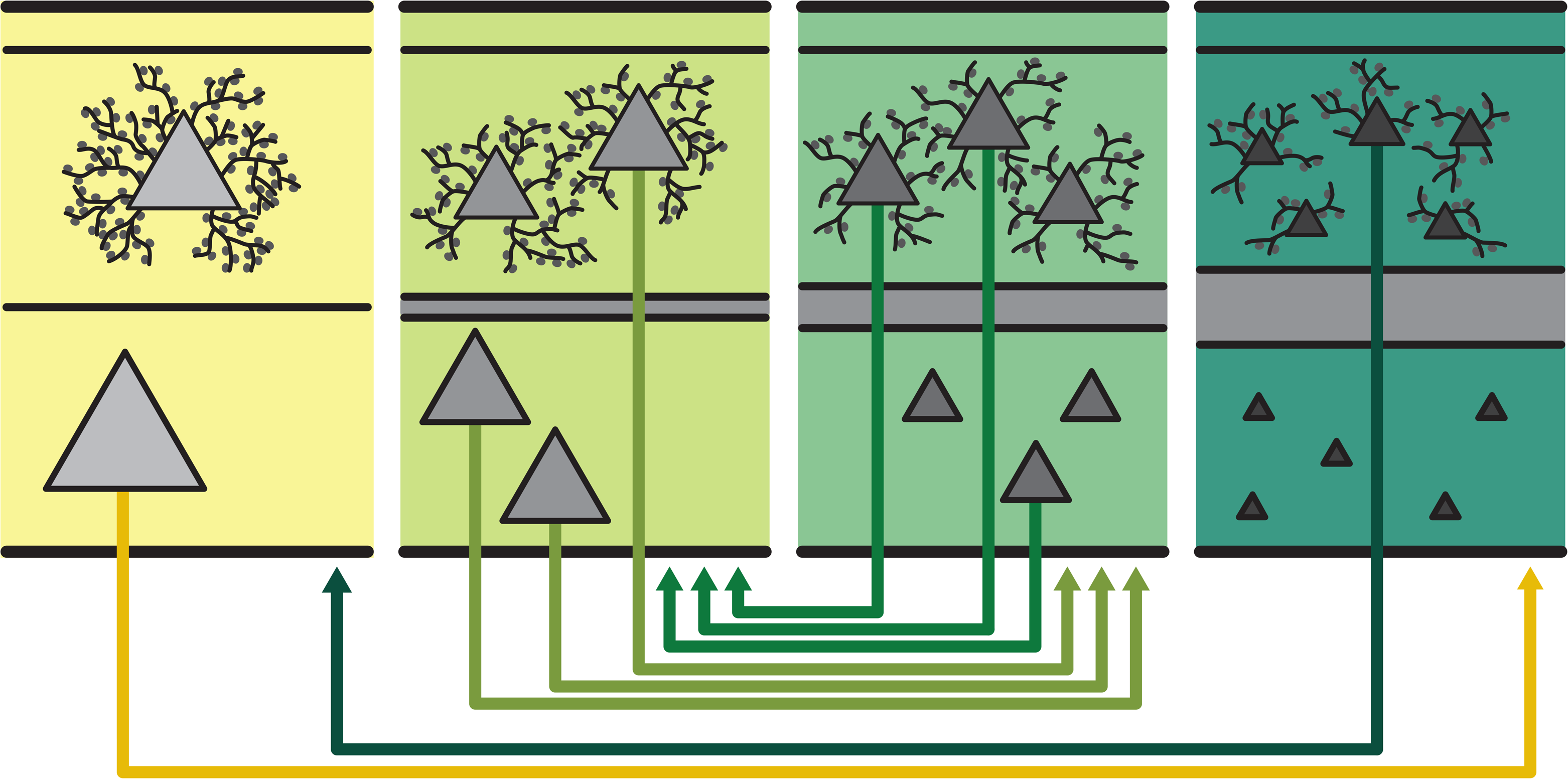
Projection patterns in context of structural variation. Both macroscopic and microscopic structural measures exhibit spatially ordered changes across the cortex. Such cortical gradients have been described for many properties of the cortical sheet (e.g., Abbie, 1940; Sanides, 1962; Zilles and Palomero-Gallagher, 2017), and are closely tied to the organization of structural and functional connections (cf. Figure 8). Here we find that less architectonically differentiated cortical areas (agranular, yellow) are characterized by lower neuron density and different morphology of layer 3 pyramidal cells than more strongly differentiated areas (eulaminate, dark green), with gradual changes across the spectrum (light green, medium green). Specifically, as shown in Table 1, higher neuron density correlates with smaller size of the soma and the dendritic tree as well as with lower total spine count and peak spine density. Laminar patterns of projection origin are indicated as observed in this report (cf. Table 5) and consistent with the architectonic type principle of cortical connectivity (Barbas, 1986; Barbas, 2015). Connections between areas of similar architectonic differentiation show a bilaminar projection origin pattern (light green to medium green, medium green to light green), while connections between areas of distinct differentiation show a skewed unilaminar projection pattern, with projections originating predominantly in the infragranular or supragranular layers (yellow to dark green, representing agranular to eulaminate projections, and dark green to yellow, representing eulaminate to agranular projections).

Moreover, neuron density was the only structural measure that was correlated with the topological measure of overall area degree, that is, the number of afferent and efferent connections of cortical areas (Figure 6, Table 6). Considering in-degree and out-degree separately revealed a strong negative correlation between out-degree and neuron density, even if all other measures were controlled for. That is, areas of weaker differentiation tended to innervate more areas than more strongly differentiated areas, which would allow the former to supply modulatory input to a large part of the cortex. For in-degree, we observed a moderate to strong correlation with both neuron density and dendritic tree size, both within the 29×29 subgraph and cortex-wide, if the measures were considered individually. These correlations were not robust enough to remain significant if all other measures were controlled for. This could be due to an actual lack of a relationship with in-degree, or added noise from the other three cellular morphological measures (which were not significantly correlated with in-degree individually) could have made a relationship indiscernible. In the latter case, areas of weaker differentiation as well as areas with larger dendritic tree sizes would tend to be targeted by more projections. This would be in line with less differentiated areas being set up for the integration of sensory inputs over a relatively large part of the cortex and meta-processing (e.g., Goldman-Rakic, 1988; Buckner and Krienen, 2013). Moreover, a larger number of incoming afferents might necessitate more dendritic space to be accommodated.

Regardless of their interpretation, the results for in-degree within the 29×29 subgraph were very similar to the results for cortex-wide in-degree. This observation corroborates a previous proposition (Ercsey-Ravasz et al., 2013) stating that the 29×29 edge-complete subgraph, whose constituent areas were widely distributed within the complete set of 91 cortical areas, is representative of the cortex-wide full network of interareal connections. Our analysis of in-degree, thus, indicates that the results for overall degree as well as out-degree, within the 29×29 subgraph, reflect genuine cortex-wide relationships between the structural measures and area degree. In summary, our analyses indicate that, while the cellular morphological measures and the area-based measure neuron density are closely related, neuron density is a more essential predictor of three of the four tested basic features of corticocortical connectivity (projection existence, laminar projection patterns and area degree) than the cellular morphological measures. Thus, our analyses unify various previous reports that related different aspects of cortical architecture to each other as well as to features of cortical connectivity. This finding is consistent with previous reports which demonstrated that neuron density provided a more characteristic ‘fingerprint’ of the cytoarchitecture of cortical areas than other architectonic measures (Dombrowski et al., 2001), as well as with reports showing that a close relation between architectonic differentiation and corticocortical connectivity could also be observed in different mammalian species such as the cat, (Hilgetag and Grant, 2010; Beul et al., 2015), the mouse (Rubinov et al., 2015; Goulas et al., 2017), and humans (van den Heuvel et al., 2015). Moreover, we demonstrated *in silico* that, given an empirically grounded relationship between the formation time of areas and their architectonic differentiation (of which neuron density is a very good surrogate measure), spatiotemporal interactions between forming cortical areas were sufficient to give rise to connectivity that conformed to the architectonic type principle (Beul et al., 2018), as it has been observed in mammalian cortico-cortical connectivity. The emergence of neuron density as a fundamental marker that relates both to structural connectivity and to further architectonic measures, as reported here, is entirely consistent with these possible mechanistic underpinnings of the architectonic type principle.

### Developmental mechanisms may regulate the covariation of architectonic measures

Systematic, joint variation of different features of cellular morphology has been observed between cortical regions within mammalian species. In primates, a higher number and higher density of spines and more complex dendritic arbors have been reported in prefrontal cortices compared to motor or sensory cortices (Elston, 2003; Elston, 2007; Elston et al., 2011a; Bianchi et al, 2013). In mouse cortex, spine density in the prelimbic and infralimbic fields is twice as high as in the remaining cortex (Ballesteros-Yáñez et al., 2010), and spine size varies across the cortex (Benavides-Piccione et al., 2002). Here, we have shown that these gradual changes in cell morphology are aligned with the overall degree of architectonic differentiation observed in cortical areas, by reporting a negative relationship between architectonic differentiation and morphological complexity. Figure 7 gives an overview of the observed relations between the five structural measures. Cortical gradients, that is, spatially ordered changes across the cortical sheet, have been described for these and multiple other macroscopic and microscopic structural measures (e.g., Abbie, 1940; Sanides, 1962; Zilles and Palomero-Gallagher, 2017), and are closely related to the organization of structural and functional connections across the cortex (see ‘Functional Implications’ below). Moreover, it has been noted that variation in both cellular architecture and neuron numbers is well aligned with developmental gradients (Charvet and Finlay, 2014; Charvet et al., 2015). This link has been corroborated by findings in the human cortex, which directly traced the systematic architectonic variation of the cortex to the timing of development (Barbas and García-Cabezas, 2016). Thus, multiple dimensions of cellular morphology appear to be tightly coupled, matching the overall degree of area differentiation as well as variation in developmental timing. Together, these observations point towards a precise orchestration of cell specification during ontogenesis, such that morphological, microscopic features of neurons and the macroscopic architectonic differentiation of an area as a whole grow attuned.

Additionally, physical self-organization may play a role in shaping the covariation of overall architectonic differentiation, determined by spatiotemporal developmental gradients, and morphological measures. Assuming that neurons and neuropil are packed into the available cortical volume as tightly as possible, approximating maximum volume packing (e.g., Chklovskii et al., 2002), the cellular morphological features would be expected to co-vary with neuron density, as is observed. In particular, a higher density of neurons would be accompanied by smaller somata, less extensive dendritic arborization, and possibly also dendrites that are less spiny.

### Functional implications

It appears that local cortical architecture and connection features of a cortical area, as well as an area’s functional role within the cortical network, are tightly interrelated (Figure 8). First, laminar projection patterns place origins and terminations in laminar microenvironments appropriate for the type of information exchange between pairs of cortical areas. As noted before, the observed anatomical distinctions between laminar patterns of projections connecting areas of varying relative differentiation likely reflect differences in information processing (e.g., Barone et al, 1995). Specifically, projections that propagate information towards more abstract and multi-modal processing regions show a different laminar composition than projections that feed back the results of information integration to areas closer to the sensory periphery, affecting behaviour by modulating information processing (reviewed in Batardière et al., 1998; Buckner and Krienen, 2013; Harris and Shepherd, 2015). These anatomical distinctions are accompanied by differences in electrophysiological signatures associated with the respective pathways (Bastos et al., 2015). Moreover, neurons in infragranular and supragranular layers have been shown to possess different physiological (Lagae et al., 1989; Nowak et al., 1995; Raiguel et al., 1995) as well as histochemical (Hof et al., 1996; 1997) characteristics. These observations are functionally relevant, since even small variations in cell-intrinsic properties can induce substantial differences in the computations performed by otherwise similar circuits (Harris and Shepherd, 2015). A large body of work on hierarchical predictive coding integrates aspects of connectivity such as intrinsic, local microcircuitry, laminar projection patterns and oscillatory signatures of pathways to explain the perception of sensory signals (e.g., Bastos et al., 2012; Friston et al., 2015; Shipp, 2016). The modulation of information processing is accomplished through precisely targeted inputs (reviewed in Larkum, 2013), so that exerting a modulatory effect does not require an accumulation of incoming connections, but can be achieved by relatively weak inputs. This situation is in contrast to driving inputs delivered by feedforward projections that originate predominantly in supragranular layers. In a comparison of feedforward and feedback projections that were similar in their absolute deviation from bilaminar projection patterns (i.e., |*N*_SG_%|), feedforward projections were reported to be stronger than feedback projections (Markov et al., 2013b), illustrating diverging requirements for driving and modulating influences. Similarly, feedforward projections terminating in middle cortical layers were shown to have larger boutons, and hence potentially stronger drive, than projections terminating outside the middle layers (Germuska et al., 2006). Moreover, the stronger, driving effect of feedforward projections is counterbalanced by their more pronounced capacity to inhibit, as reported by D’Souza and colleagues (2016), who have shown that the fraction of inhibitory targets is larger for feedforward than feedback projections. Connection features thus correspond well to the functional roles of connected areas. Second, in a complementary way, the intrinsic processing capabilities of cortical areas are, to a large extent, determined by local characteristics of cellular morphology. Cellular morphological properties and the functional roles of cortical areas, thus, also appear attuned to each other. Finally, the relation between cortical architecture and connection features completes the three interactive aspects. The observations integrated into the architectonic type principle extensively describe this interrelation. Moreover, the dependence of cortical connections on relative architectonic differentiation formalized in the architectonic type principle affords deeper insight into multiple aspects of the organization of cortical connectivity (reviewed in Barbas, 2015; Hilgetag et al., 2016). For example, the arrangement of areas by relative differentiation is consistent with onset latencies in the visual system (Petroni et al., 2001) and an ordering of areas inferred from their functional interactions in different frequency bands (Bastos et al., 2015). Laminar projection patterns indicate that areas of weak differentiation (limbic cortices) feed back information to more strongly differentiated areas (Barbas, 1986; Barbas and Rempel-Clower, 1997), placing them in a central position in the cortical network. This observation is consistent with their involvement in the default mode network (Raichle, 2011) as well as a suggested link to functional conscious access (Dehaene et al., 1998; 2011). The architectonic type principle also pertains to disruptions of specific pathways in diseases, for example autism (Zikopoulos and Barbas, 2010; 2013). In summary, the strong correlation between the laminar origins of interareal projections and relative architectonic differentiation is closely intertwined with the functional relevance of a given projection. It reflects the seamless integration of the functional interplay of areas with local morphological properties and their associated intrinsic processing capabilities.

**Figure 8.**
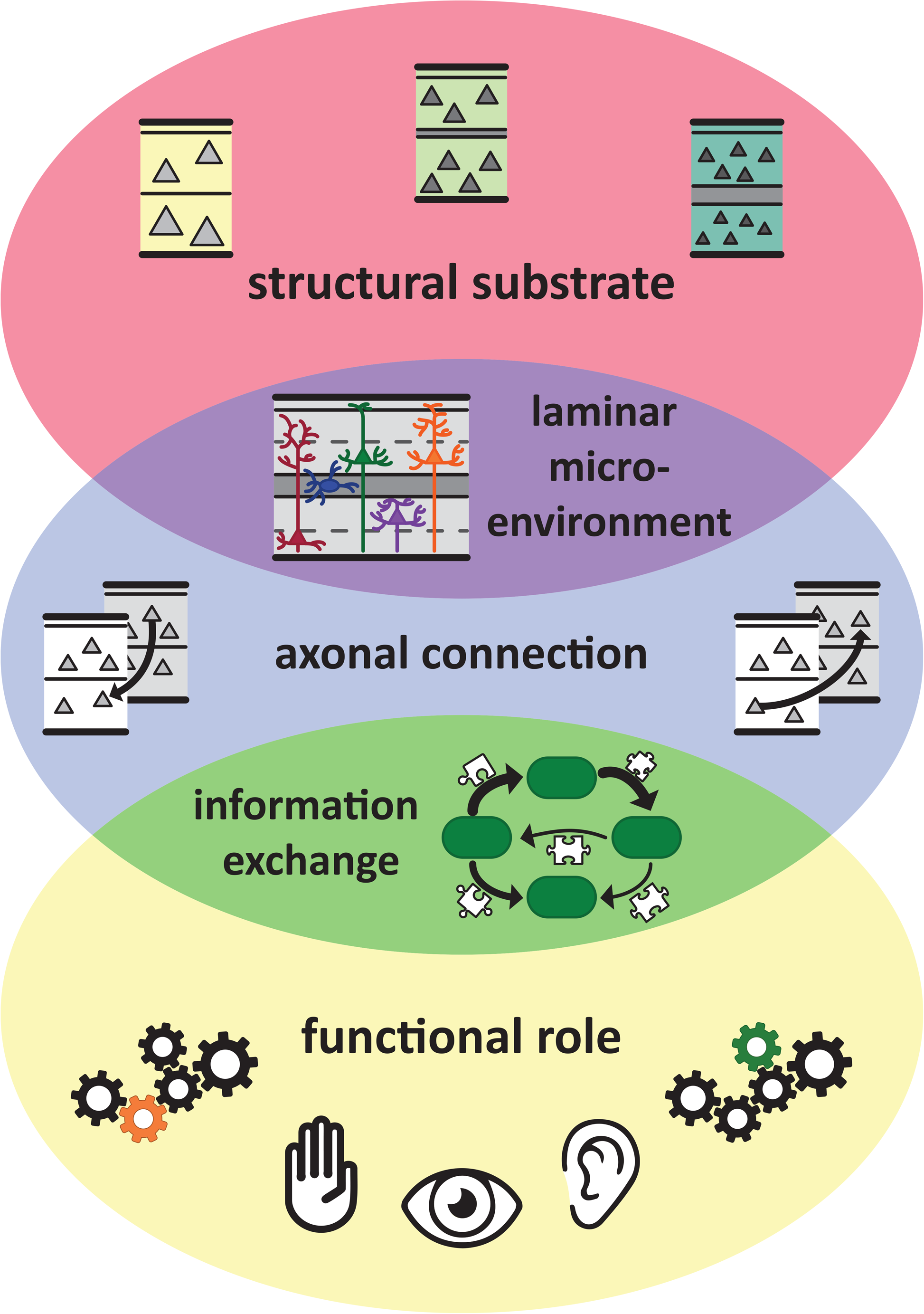
Structure and function of brain areas and connections are interlinked. Connections create function of brain areas and functional interactions among brain areas from the structural substrate of the brain, the cortical sheet. In particular, areas are linked through connections which have a laminar composition that is appropriate for the laminar microenvironment within the respective areas and the type of information exchange between these areas. Thus, local cortical architecture, the connection features of a cortical area, and an area’s functional role within the cortical network are tightly intertwined. See ‘Functional Implications’ for an elaboration of the links between these three aspects.

### Limitations of the explanatory power of architectonic factors

The present results show that a characteristic indicator of the overall degree of architectonic differentiation, neuron density, is well suited for accounting for structural connections within the global, macro-scale cortical connectome of the primate. In contrast, more fine-grained structural aspects of the cortex, such as the cellular morphological features considered here, convey less information on the connectional features of areas, appearing as derivate properties determined mostly by overall regional differentiation. However, it should be noted, that the considered morphological measures were solely acquired in supragranular cortical layer 3 and characterize only pyramidal neurons. Hence, these measures were not designed to comprehensively capture the intrinsic architectonic organization of cortical areas. Considering such inherent differences between the measures, it is plausible that neuron density, as an overall characterization of area architecture, correlates better with the areas’ macroscopic connectivity properties, as found. If a more detailed characterization of cellular morphology was available, for example through equivalent morphological measures obtained from the infragranular layers, the morphology might be captured by a summary measure (e.g., ratios across different laminar compartments) which could be used to characterize overall cortical architecture. Such a more detailed characterization of cellular morphology might then correlate with macroscopic properties of the connectome as well as neuron density. Moreover, it will be interesting to see how such findings might vary across the spectrum of mammalian cortical organization, considering that the degree to which architectonic gradients exist within the cortex of a species is variable across mammals.

### Conclusions

Cortical architecture has been shown to relate to fundamental aspects of the organization of corticocortical connections (Barbas, 1986; Barbas and Rempel-Clower, 1997; Barbas et al., 2005; Medalla and Barbas, 2006; Hilgetag and Grant, 2010; Barbas, 2015; Beul et al., 2015; Hilgetag et al., 2016; Beul et al., 2017) and appears integral for understanding cortical connectivity. However, not all structural measures are equally informative on connectivity, as we have shown in conjoint analyses of multiple macro-and microscopic properties here, as well as in previous reports (Dombrowski et al., 2001; Hilgetag et al., 2016; Beul et al., 2017). We found that neuron density, a basic and classic (Brodmann, 1909) macroscopic indicator of overall architectonic differentiation of cortical areas, more consistently related to multiple features of the macaque connectome than four microscopic morphological measures. These cellular measures, moreover, were themselves closely related to neuron density. Thus, it can be speculated whether these microscopic properties are developmentally attuned to overall architectonic differentiation of the cortex. Such an alignment might result from neurodevelopmental mechanisms combining genetic determination of regionally specific gradients with processes of physical self-organization, such as maximum volume packing, resulting in trade-offs between cell density and cell size as well as cellular complexity. Neuron density, thus, is a fundamental feature which links the macroscale and microscale architectonic and connectional organization of cortical areas and allows integrating the overall structural differentiation of areas with features of cellular morphology as well as with the existence and laminar characteristics of corticocortical connections. This insight may be a significant step towards the development of advanced across-scales models of cortical organization and function.

## Acknowledgements

We thank Alexandros Goulas for helpful comments on the manuscript. SFB and CCH were supported by DFG Collaborative Research Center Grant SFB 936/A1. Address correspondence to Claus C. Hilgetag, Institute of Computational Neuroscience, University Medical Center Eppendorf, Martinistr. 52, 20246 Hamburg, Germany.

## Supplementary Data

**Supplementary Table 1.**
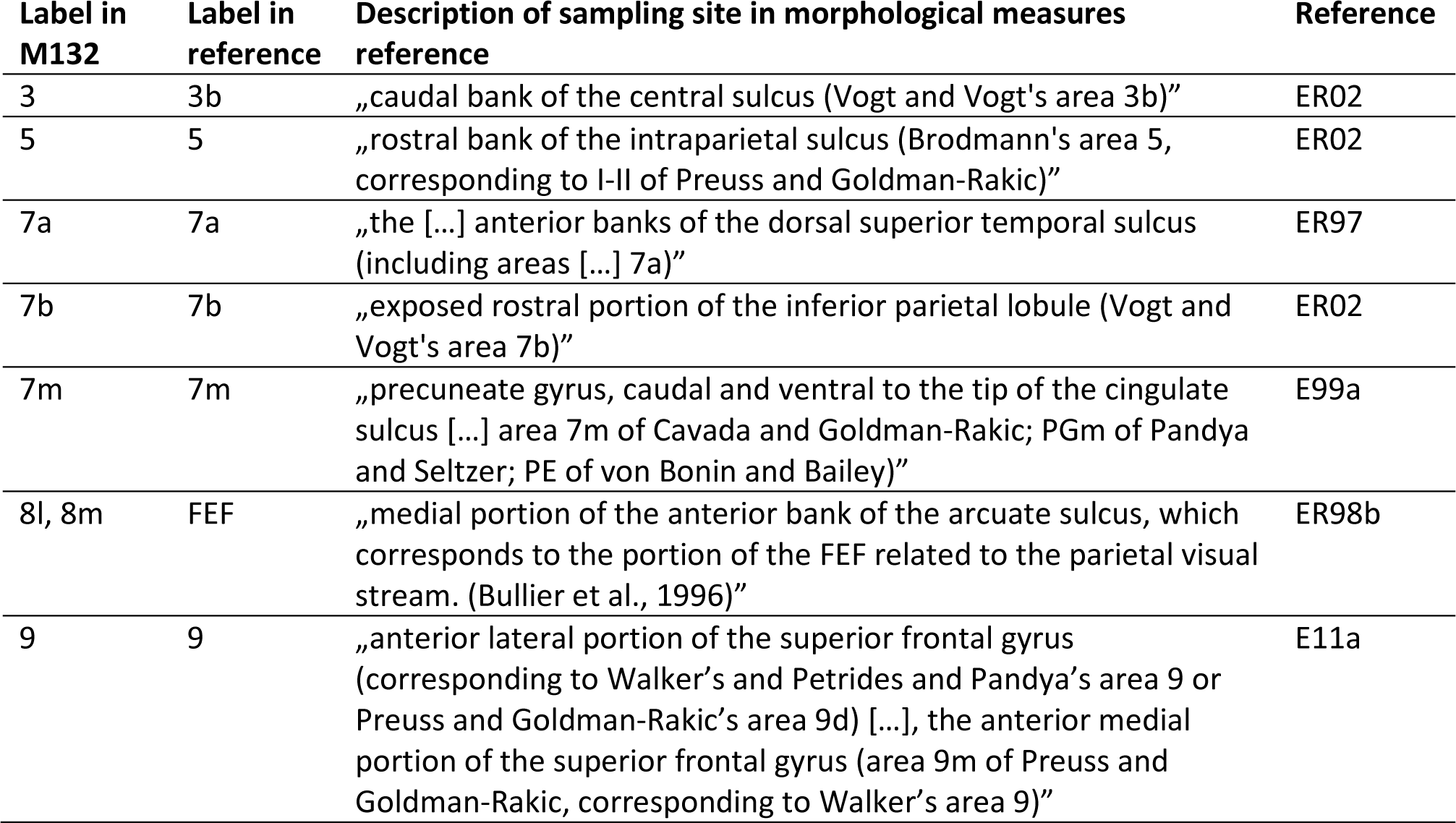
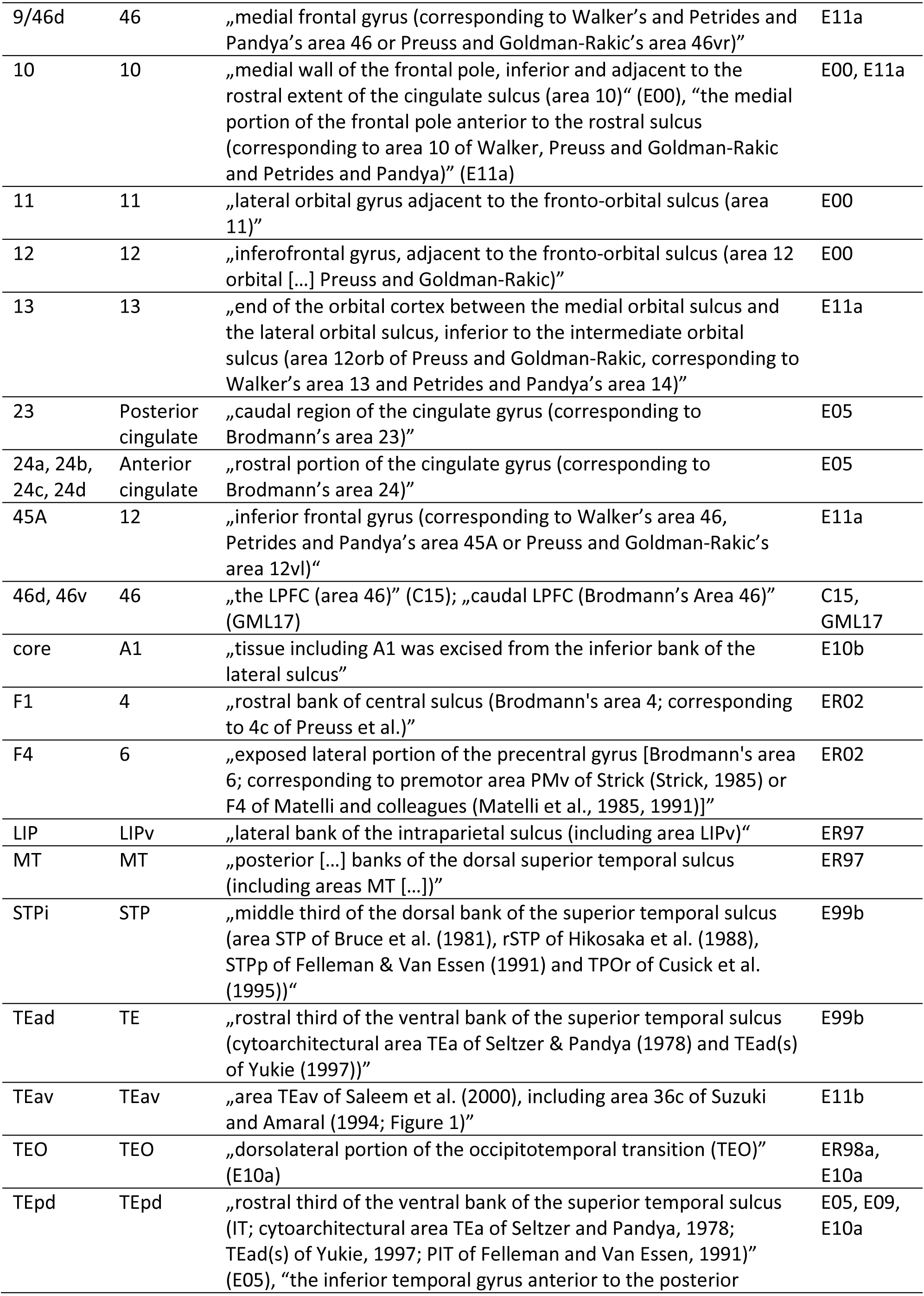
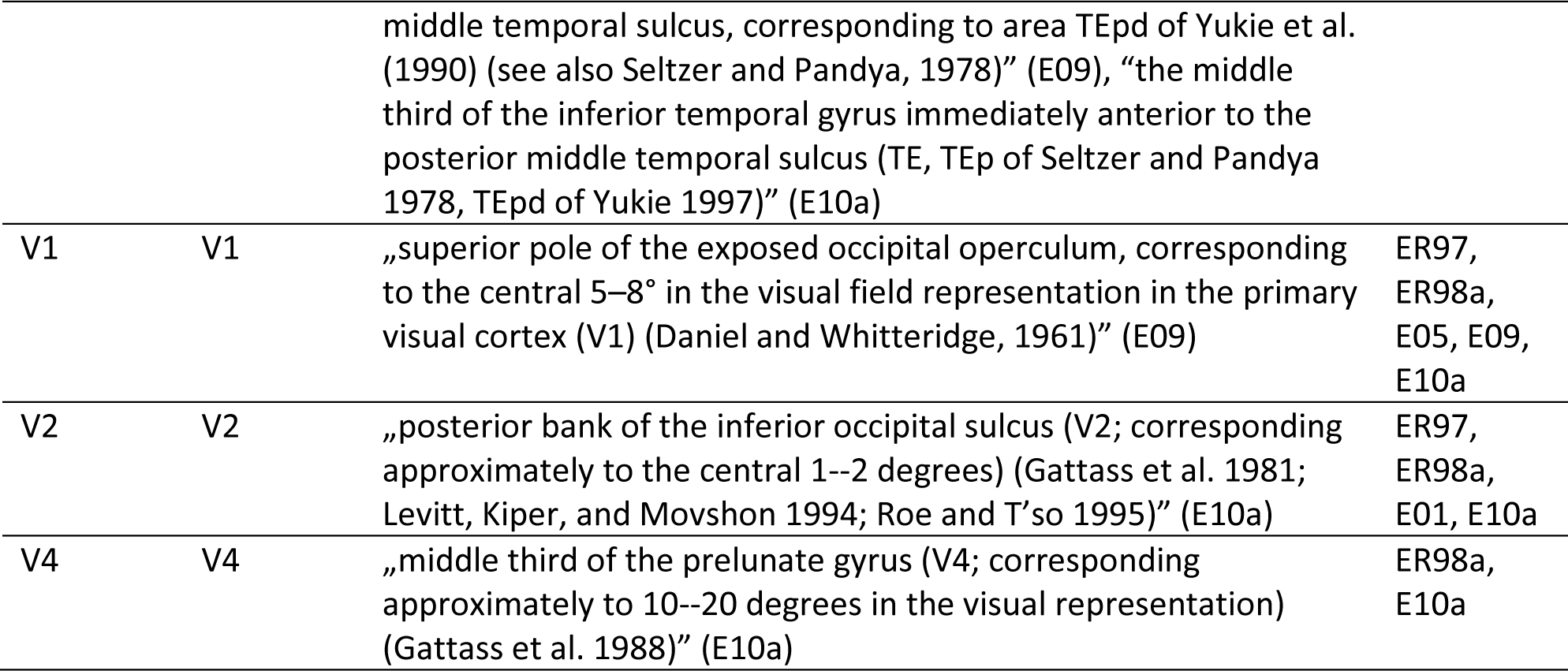
Correspondence of cortical areas across parcellations. Connectivity data were published in the M132 parcellation (Markov et al., 2014), while the morphological measures were published in reports using alternative parcellations of the macaque cortex. This table shows how areas were matched across parcellations and indicates which references the morphological measures were collated from. See main text for references. C15, Coskren et al. 2015; ER97, Elston and Rosa 1997; ER98a, Elston and Rosa 1998a; ER98b, Elston and Rosa 1998b; E99a, Elston et al., 1999a; E99b, Elston et al., 1999b; E00, Elston 2000; E01, Elston et al. 2001; ER02, Elston and Rockland 2002; E05, Elston et al. 2005; E09, Elston et al. 2009; E10a, Elston et al. 2010a; E10b, Elston et al. 2010b; E11a, Elston et al. 2011a; E11b, Elston et al. 2011b, GML17, Gilman et al. 2017.

| Label in M132 | Label in reference | Description of sampling site in morphological measures reference | Reference |
| --- | --- | --- | --- |
| 3 | 3b | „caudal bank of the central sulcus (Vogt and Vogt's area 3b)“ | ER02 |
| 5 | 5 | „rostral bank of the intraparietal sulcus (Brodmann's area 5, corresponding to I-II of Preuss and Goldman-Rakic)“ | ER02 |
| 7a | 7a | „the [...] anterior banks of the dorsal superior temporal sulcus (including areas [...] 7a)“ | ER97 |
| 7b | 7b | „exposed rostral portion of the inferior parietal lobule (Vogt and Vogt's area 7b)“ | ER02 |
| 7m | 7m | „precuneate gyrus, caudal and ventral to the tip of the cingulate sulcus [...] area 7m of Cavada and Goldman-Rakic; PGm of Pandya and Seltzer; PE of von Bonin and Bailey)“ | E99a |
| 8l, 8m | FEF | „medial portion of the anterior bank of the arcuate sulcus, which corresponds to the portion of the FEF related to the parietal visual stream. (Bullier et al., 1996)“ | ER98b |
| 9 | 9 | „anterior lateral portion of the superior frontal gyrus (corresponding to Walker's and Petrides and Pandya's area 9 or Preuss and Goldman-Rakic's area 9d) [...], the anterior medial portion of the superior frontal gyrus (area 9m of Preuss and Goldman-Rakic, corresponding to Walker's area 9)“ | E11a |
| 9/46d | 46 | „medial frontal gyrus (corresponding to Walker’s and Petrides and Pandya’s area 46 or Preuss and Goldman-Rakic’s area 46vr)” | E11a |
| 10 | 10 | „medial wall of the frontal pole, inferior and adjacent to the rostral extent of the cingulate sulcus (area 10)” (E00), “the medial portion of the frontal pole anterior to the rostral sulcus (corresponding to area 10 of Walker, Preuss and Goldman-Rakic and Petrides and Pandya)” (E11a) | E00, E11a |
| 11 | 11 | „lateral orbital gyrus adjacent to the fronto-orbital sulcus (area 11)” | E00 |
| 12 | 12 | „inferofrontal gyrus, adjacent to the fronto-orbital sulcus (area 12 orbital [...]) Preuss and Goldman-Rakic)” | E00 |
| 13 | 13 | „end of the orbital cortex between the medial orbital sulcus and the lateral orbital sulcus, inferior to the intermediate orbital sulcus (area 12orb of Preuss and Goldman-Rakic, corresponding to Walker’s area 13 and Petrides and Pandya’s area 14)” | E11a |
| 23 | Posterior cingulate | „caudal region of the cingulate gyrus (corresponding to Brodmann’s area 23)” | E05 |
| 24a, 24b, 24c, 24d | Anterior cingulate | „rostral portion of the cingulate gyrus (corresponding to Brodmann’s area 24)” | E05 |
| 45A | 12 | „inferior frontal gyrus (corresponding to Walker’s area 46, Petrides and Pandya’s area 45A or Preuss and Goldman-Rakic’s area 12vl)” | E11a |
| 46d, 46v | 46 | „the LPFC (area 46)” (C15); „caudal LPFC (Brodmann’s Area 46)” (GML17) | C15, GML17 |
| core | A1 | „tissue including A1 was excised from the inferior bank of the lateral sulcus” | E10b |
| F1 | 4 | „rostral bank of central sulcus (Brodmann’s area 4; corresponding to 4c of Preuss et al.)” | ER02 |
| F4 | 6 | „exposed lateral portion of the precentral gyrus [Brodmann’s area 6; corresponding to premotor area PMv of Strick (Strick, 1985) or F4 of Matelli and colleagues (Matelli et al., 1985, 1991)]” | ER02 |
| LIP | LIPv | „lateral bank of the intraparietal sulcus (including area LIPv)” | ER97 |
| MT | MT | „posterior [...] banks of the dorsal superior temporal sulcus (including areas MT [...])” | ER97 |
| STPi | STP | „middle third of the dorsal bank of the superior temporal sulcus (area STP of Bruce et al. (1981), rSTP of Hikosaka et al. (1988), STPp of Felleman & Van Essen (1991) and TPOr of Cusick et al. (1995))” | E99b |
| TEad | TE | „rostral third of the ventral bank of the superior temporal sulcus (cytoarchitectural area TEa of Seltzer & Pandya (1978) and TEad(s) of Yukie (1997))” | E99b |
| TEav | TEav | „area TEav of Saleem et al. (2000), including area 36c of Suzuki and Amaral (1994; Figure 1)” | E11b |
| TEO | TEO | „dorsolateral portion of the occipitotemporal transition (TEO)” (E10a) | ER98a, E10a |
| TEpd | TEpd | „rostral third of the ventral bank of the superior temporal sulcus (IT; cytoarchitectural area TEa of Seltzer and Pandya, 1978; TEad(s) of Yukie, 1997; PIT of Felleman and Van Essen, 1991)” (E05), “the inferior temporal gyrus anterior to the posterior middle temporal sulcus, corresponding to area TEpd of Yukie et al. (1990) (see also Seltzer and Pandya, 1978)” (E09), “the middle third of the inferior temporal gyrus immediately anterior to the posterior middle temporal sulcus (TE, TEp of Seltzer and Pandya 1978, TEpd of Yukie 1997)” (E10a) | E05, E09, E10a |
| V1 | V1 | „superior pole of the exposed occipital operculum, corresponding to the central 5–8° in the visual field representation in the primary visual cortex (V1) (Daniel and Whitteridge, 1961)” (E09) | ER97, ER98a, E05, E09, E10a |
| V2 | V2 | „posterior bank of the inferior occipital sulcus (V2; corresponding approximately to the central 1–2 degrees) (Gattass et al. 1981; Levitt, Kiper, and Movshon 1994; Roe and T’so 1995)” (E10a) | ER97, ER98a, E01, E10a |
| V4 | V4 | „middle third of the prelunate gyrus (V4; corresponding approximately to 10–20 degrees in the visual representation) (Gattass et al. 1988)” (E10a) | ER98a, E10a |

Supplementary Tables 2 to 5

All analyses reported in supplementary tables 2 to 5 were performed in the same way as the corresponding main analyses (see Tables 1 to 6), but on a reduced connectivity data set, from which all projections with less than five constituent axons were excluded (see Results). Results corresponding to Table 1 are not presented here, because the structural measures did not change through applying a cutoff to the connectivity data set. Results corresponding to Table 5 are not presented here, because we applied a cutoff of 20 constituent axons to the connectivity data set in the main analyses of laminar projection patterns and no further connections were excluded by the lower threshold of five constituent axons.

**Supplementary Table 2.** Structural measures in connected and unconnected pairs of areas in reduced data set. Absolute values of relative structural measures are averaged across area pairs without (absent) and with (present) a linking projection. T-statistics, p-values and effect size *r* are results of two-tailed independent samples t-tests comparing the two respective distributions for equal means.

|  | Absent projections |  | Present projections |  | t-statistic | p-value | r |
| --- | --- | --- | --- | --- | --- | --- | --- |
|  | Mean | Group size | Mean | Group size |  |  |  |
| Area-based measures |  |  |  |  |  |  |  |
| \|log-ratio<sub>density</sub>\| | 0.67 | 402 | 0.33 | 641 | t<sub>(1041)</sub> = 14.5 | <.001 | 0.41 |
| geodesic distance [mm] | 30.9 | 995 | 22.6 | 1410 | t<sub>(2403)</sub> = 22.0 | <.001 | 0.41 |
| Cellular morphological measures |  |  |  |  |  |  |  |
| \|Δ<sub>soma size</sub>\| [μm<sup>2</sup>] | 71.7 | 190 | 58.8 | 288 | t<sub>(476)</sub> = 2.7 | 0.007 | 0.12 |
| \|Δ<sub>spine count</sub>\| [#] | 3525 | 205 | 2687 | 290 | t<sub>(493)</sub> = 4.3 | <.001 | 0.19 |
| \|Δ<sub>spine density</sub>\| [# / 10μm] | 8.9 | 205 | 7.3 | 290 | t<sub>(493)</sub> = 3.1 | 0.002 | 0.14 |
| \|Δ<sub>tree size</sub>\| [μm<sup>2</sup>] | 48420 | 220 | 37191 | 323 | t<sub>(541)</sub> = 4.0 | <.001 | 0.17 |

**Supplementary Table 3.**
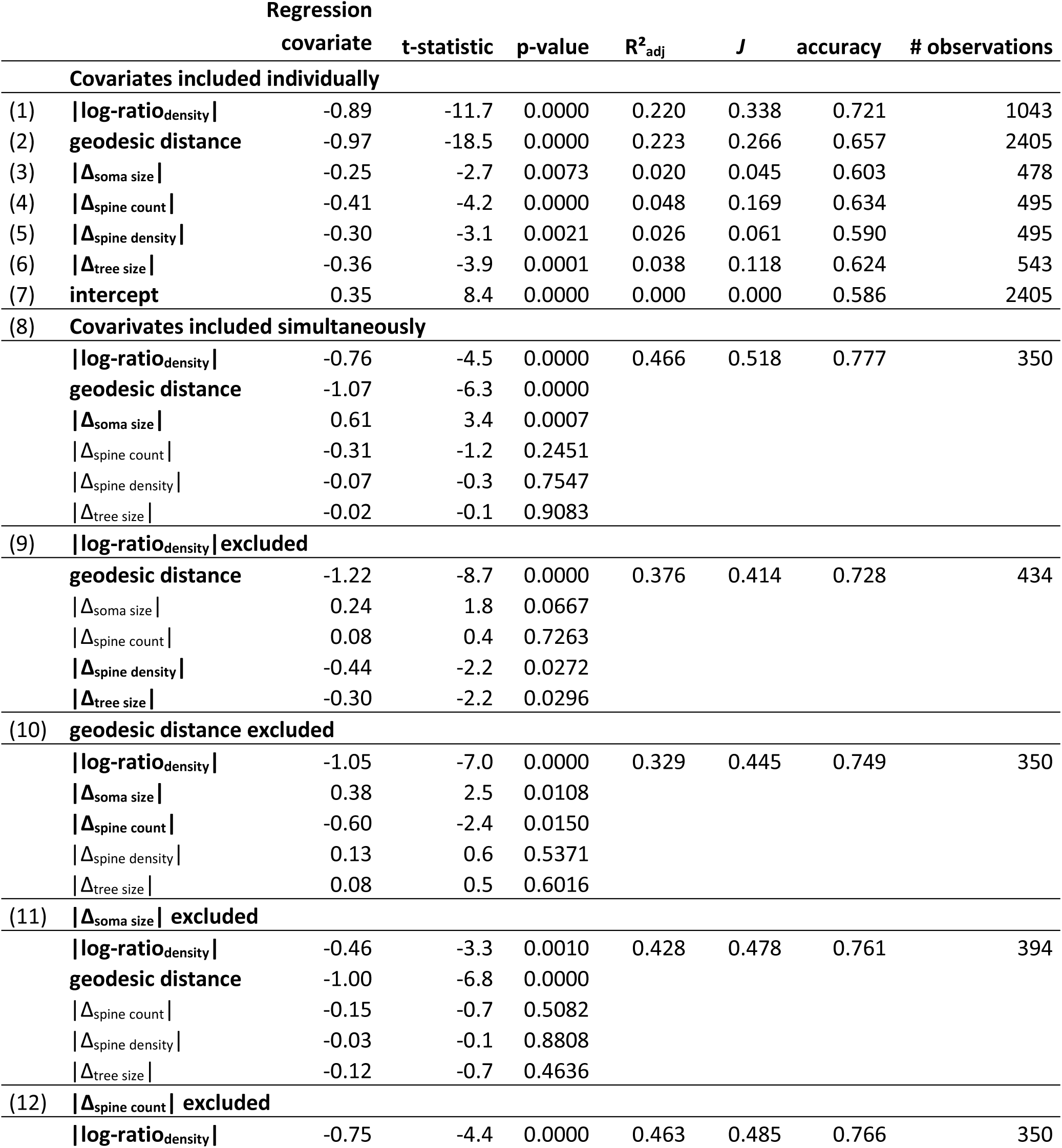
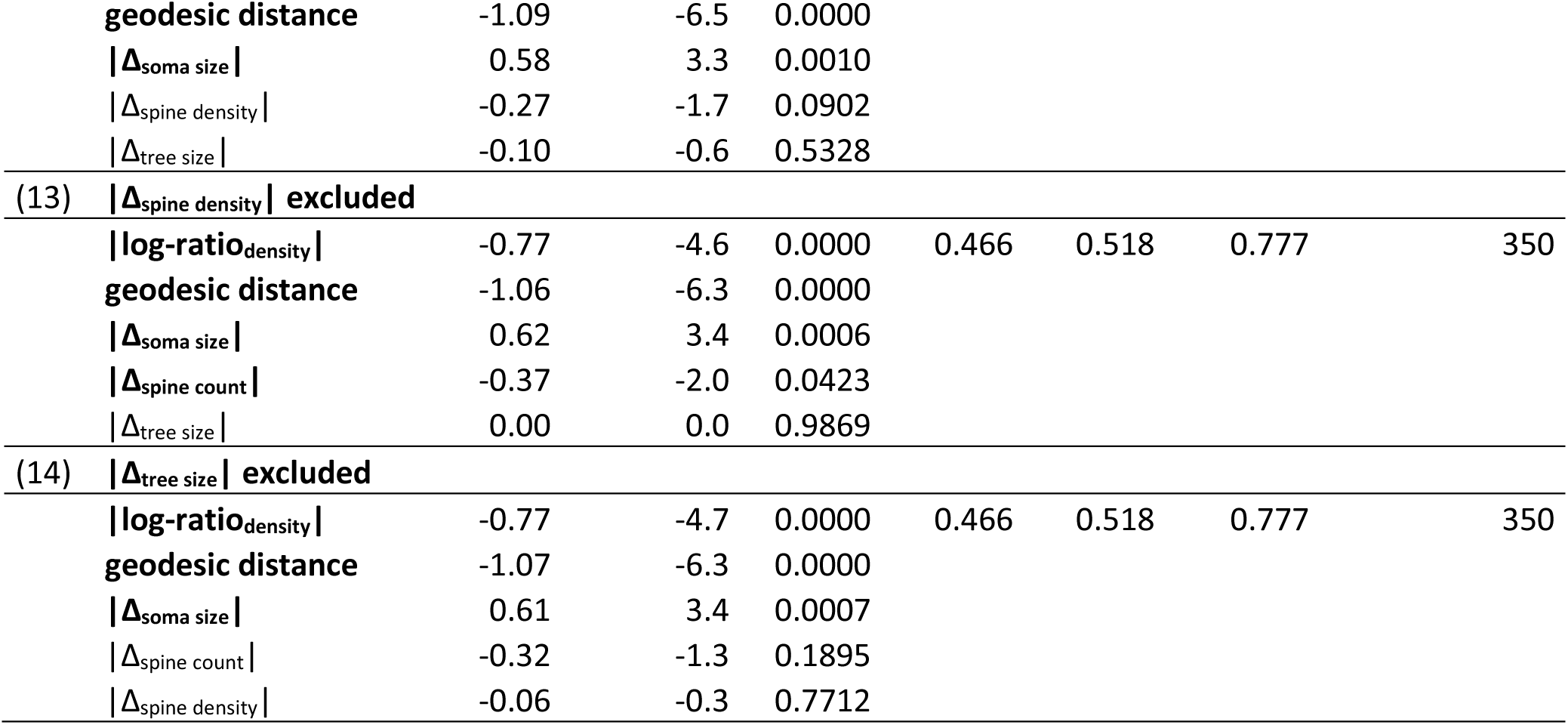
Classification of connection existence by logistic regression in reduced data set. We performed binary logistic regression analyses (enumerated in brackets), each including a different set of the structural measures as covariates and connectivity (grouped into ‘absent’ and ‘present’ connections) as the dependent variable. Bold-faced covariates significantly contributed to classification performance as indicated by the p-value. Across all regression analyses, absolute neuron density ratio, geodesic distance and soma size difference consistently emerged as meaningful predictors.

**Supplementary Table 4.** Correlation between projection strength and structural measures in reduced data set. Pearson correlation coefficients and associated p-values for correlations between projection strength, expressed either as ln(FLNe) or as ranked strengths, and absolute values of relative structural measures or ranked absolute values of relative structural measures. Correlations were assessed both for each measure independently (individual correlation) and while accounting for all other five measures (partial correlation).

| **FLNe** | **Individual correlation** |  | **Partial correlation** |  |
| --- | --- | --- | --- | --- |
|  | r | p-value | r | p-value |
| \|log-ratio<sub>density</sub>\| | -0.19 | <0.0001 | -0.05 | 0.4487 |
| geodesic distance | -0.47 | <0.0001 | -0.51 | <0.0001 |
| \|$\Delta_{\text{soma size}}$\| | -0.11 | 0.0516 | 0.09 | 0.1965 |
| \|$\Delta_{\text{spine count}}$\| | -0.17 | 0.0028 | 0.00 | 0.9467 |
| \|$\Delta_{\text{spine density}}$\| | -0.12 | 0.0468 | -0.13 | 0.0610 |
| \|$\Delta_{\text{tree size}}$\| | -0.13 | 0.0197 | -0.19 | 0.0056 |
| **ranked strength** | $\rho$ | p-value | $\rho$ | p-value |
| ranked \|log-ratio<sub>density</sub>\| | 0.20 | <0.0001 | 0.13 | 0.0690 |
| ranked geodesic distance | 0.44 | <0.0001 | 0.46 | <0.0001 |
| ranked \|$\Delta_{\text{soma size}}$\| | 0.15 | 0.0116 | -0.14 | 0.0528 |
| ranked \|$\Delta_{\text{spine count}}$\| | 0.12 | 0.0372 | -0.02 | 0.7246 |
| ranked \|$\Delta_{\text{spine density}}$\| | 0.09 | 0.1112 | 0.12 | 0.0836 |
| ranked \|$\Delta_{\text{tree size}}$\| | 0.15 | 0.0053 | 0.20 | 0.0052 |

**Supplementary Table 5.**
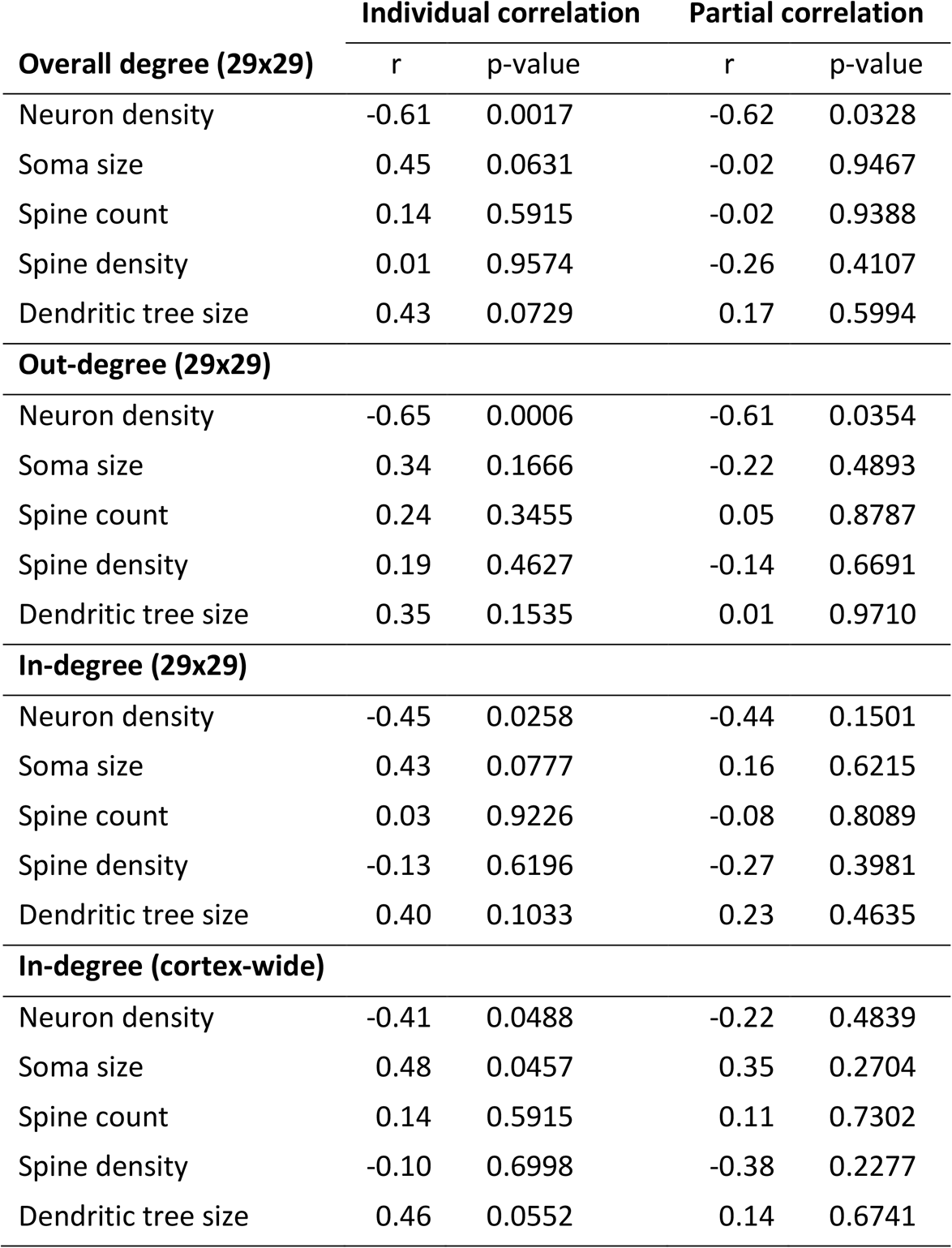
Correlation between area degree and structural measures in reduced data set. Pearson correlation coefficients and associated p-values for correlations between the structural measures for each area and overall area degree (total number of maintained connections), out-degree, in-degree or cortex-wide in-degree. Correlations were assessed both for each measure independently (individual correlation) and while accounting for the other five measures (partial correlation). Geodesic distance could not be included because it is a relational property not defined for individual areas. Bonferroni correction for multiple tests results in an adjusted significance threshold of α_adj_ = 0.05/5 = 0.01 for the individual correlations.

